# Kinesin-3 and kinesin-1 motors direct basement membrane protein secretion to a basal sub-region of the basolateral plasma membrane in epithelial cells

**DOI:** 10.1101/2021.01.31.429062

**Authors:** Allison L. Zajac, Sally Horne-Badovinac

## Abstract

Basement membranes (BMs) are sheet-like extracellular matrices that line the basal surfaces of all epithelia. Since BM proteins form networks, they likely need to be secreted near the basal surface. However, the location of their secretion site and how it is selected are unknown. Working in the *Drosophila* follicular epithelium, we identified two kinesins essential for normal BM formation. Our data suggest the two kinesins work together to transport Rab10+ BM protein-filled secretory vesicles towards the basal surface along the polarized microtubule array common to epithelia. This kinesin transport biases BM protein secretion basally. When kinesins are depleted, BM proteins are mis-secreted to more apical regions of the lateral membrane, creating ectopic BM protein networks between cells that disrupt cell movements and tissue architecture. These results introduce a new transport step in the BM protein secretion pathway and highlight the importance of controlling the sub-cellular exocytic site of network-forming proteins.

**Highlights:**

- A kinesin-3 and a kinesin-1 are required for normal basement membrane (BM) assembly
- Kinesins move Rab10+ BM secretory vesicles basally on polarized microtubule arrays
- Transport biases BM exocytosis to basal subregions of the basolateral membrane
- Loss of kinesins creates ectopic BM networks that disrupt tissue architecture

## INTRODUCTION

The basement membrane (BM) is a sheet-like extracellular matrix present in most organs in the body that plays essential roles in tissue development and physiology (Jayadev and Sherwood, 2017; Ramos-Lewis and Page-McCaw, 2018). BMs provide attachment sites for cells, are a reservoir of growth factors, and provide mechanical support to shape tissues. The main structural components of the BM are type IV collagen (Col IV), laminin, heparan sulfate proteoglycans like perlecan, and nidogen/entactin. Many other proteins are found in BMs, and variations in both protein composition and structure allow BMs to carry out important tissue-specific functions like filtering blood in the kidney (Miner, 2012), forming the transparent high refractive index lens capsule in the eye (Danysh and Duncan, 2009), and mechanically protecting muscle fibers (Sanes, 2003). Despite the ubiquity of BMs and their many essential functions, we know little about how BM proteins are secreted, and ultimately assembled, in the correct place within a tissue.

In this study, we investigate how a single sheet of BM is built at the basal surface of an epithelium. BM proteins can be secreted from other tissues and/or produced by the epithelial cells themselves; we focus on epithelial cell-produced BMs. Epithelial cells generate and maintain their polarized membrane domains in part through sorting newly made proteins into apically- or basolaterally-directed secretory pathways (Rodriguez-Boulan and Macara, 2014). Basolateral proteins are generally thought to be secreted through an apical region of the lateral membrane where Par-3 acts as a receptor for the exocyst vesicle tethering complex (Ahmed and Macara, 2017; Kreitzer et al., 2003; Polishchuk et al., 2004). However, BM proteins are large, network-forming proteins, suggesting their secretion site may need to be closer to the basal surface to prevent the assembly of ectopic networks. The secretion site for BM proteins in any epithelium is unknown.

Much of what is known about the polarized secretion of BM proteins comes from studies of the follicular epithelium of *Drosophila*. This somatic epithelium surrounds a cluster of germ cells to form an ovarian follicle (egg chamber) that will produce one egg (Figure 1A) (Horne-Badovinac and Bilder, 2005). The follicular epithelial cells (follicle cells) make their own BM proteins and build a BM on the egg chamber’s outer surface (Figure 1A-C). These features allow high-resolution, live imaging of intracellular BM protein trafficking and BM assembly in an intact, developing tissue. Moreover, the powerful genetic approaches in this system have allowed our lab and others to begin to unravel the molecular logic regulating polarized BM protein secretion, including identification of two small GTPases, Rab10 and Rab8, that are required to sort BM proteins into a basally-directed trafficking route (Denef et al., 2008; Devergne et al., 2014, 2017; Isabella and Horne-Badovinac, 2016; Lerner et al., 2013).

**Figure 1.**
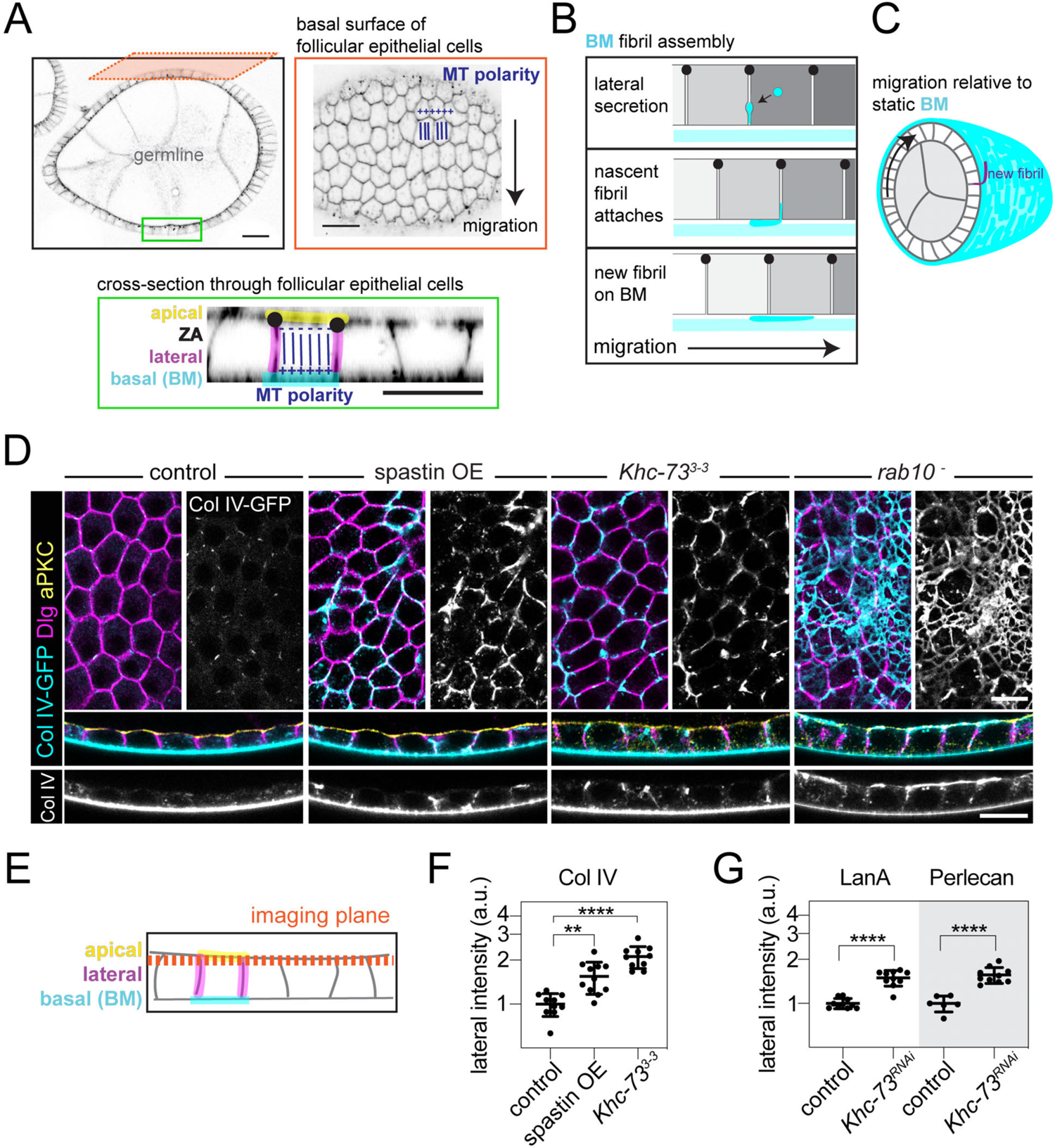
Khc-73 and MTs are required for polarized BM protein secretion. A. Images of an egg chamber with cell edges marked with CellMask. The green box highlights follicle cell polarity and MT polarity (ZA, zonula adherens). The orange box highlights a plane along the basal surface with the polarity of the basal MT array indicated relative to the direction of cell migration. B. Illustration of how BM fibrils are formed by lateral secretion, and then drawn out onto the BM during migration. C. Illustration of a transverse section through an egg chamber showing follicle cells collectively migrating along the BM, causing all cells to rotate relative to the static BM while laying down BM fibrils (purple highlight). D. Images of ectopic Col IV-GFP in epithelia overexpressing spastin or mutant for Khc-733-3 or rab10-. Top panels are planes through the lateral domains that capture some of the apical surface due to tissue curvature, as diagrammed in (E). Bottom panels are cross sections. Anti-Dlg marks lateral domains. Anti-aPKC marks apical domains (only shown in cross sections). E. Illustration of imaging planes in (D). F. Quantification of increased Col IV-GFP at lateral cell edges from (D) in epithelia overexpressing spastin or mutant for Khc-733-3. Ordinary one-way ANOVA with Dunnett’s multiple comparisons test, **p<0.01, ****p < 0.0001. In the order on graph, n= 10,11,10 egg chambers. G. Quantification of increased LanA and Perlecan accumulation at lateral cell edges in control and Khc-73RNAi epithelia. Unpaired t test, ****p<0.0001. In the order on graph, n= 10,10,6,10 egg chambers. Stage 8 egg chambers. Data represent mean ± SD plotted on a log scale. Scale bars, (A) 20 µm, all others 10 µm. See also Figure S1 and Movie 1.

Studies of the follicle cells have also shown that the location where new BM proteins are secreted can affect the architecture of the resulting BM network. During egg chamber development, the follicular BM becomes mechanically anisotropic. The BM is stiffer around the center of the egg chamber and softer at the ends (Crest et al., 2017), allowing it to act as a “molecular corset” (Gutzeit, 1991) that constrains growth in one direction, promoting tissue elongation as the egg chamber develops (Figure S1A) (Crest et al., 2017; Gutzeit, 1991). One important contributor to the stiffness of this BM is the organization of a subset of BM proteins into an aligned array of fibril-like structures (Chlasta et al., 2017; Crest et al., 2017), which are created by coupling BM protein secretion with the collective migration of follicle cells along the BM (Figures 1B-C and Movie 1) (Isabella and Horne-Badovinac, 2016). During this process, some BM protein secretion is targeted to the lateral membrane, which allows this population of BM proteins to form nascent fibrils in the space between the cells (Figure 1B). Attachment of a nascent fibril to the BM then causes it to be drawn out onto the BM sheet as cells migrate away from this anchor point (Figures 1B and 1C). From our work on BM fibril formation, we proposed that BM protein secretion may be restricted to basal regions of the lateral membrane and basal surface. However, where BM protein exocytosis occurs in the follicle cells and the mechanisms that direct secretion to this site remain to be determined.

One potential mechanism to bias the site of BM protein secretion to basal regions of the plasma membrane is suggested by the organization of MTs in epithelial cells. These cells have a polarized array of MTs whose minus ends are anchored apically and plus ends grow toward the basal surface (Figure 1A) (Sanchez and Feldman, 2017). MTs in mammalian epithelial cells are required for polarized BM protein secretion (Almeida and Stow, 1991; Boll et al., 1991). Since MT plus ends are enriched basally, transport of BM protein-filled secretory vesicles by plus end-directed kinesin motors could bias secretion to regions of the plasma membrane near the BM. Vesicular transport often requires multiple different kinesins, whose distinct properties allow them to collectively navigate the crowded cytoplasm and carry their cargo to the correct destination in the cell (Burute and Kapitein, 2019; Hancock, 2014). However, the mechanistic role MTs play in BM protein secretion, and whether kinesins mediate this process, are unknown.

Here we show that kinesin-based transport biases the site of BM protein secretion to the basal-most regions of the basolateral plasma membrane. Our data support a model in which the combined activity of the kinesin-3 motor, Khc-73, and the kinesin-1 motor, Khc, are needed to transport Rab10+ BM protein-filled vesicles along the polarized MT array to the correct secretion site near the basal surface. When this transport is reduced, some BM proteins are mis-secreted through more apical regions of the lateral membrane. This leads to the formation of an ectopic BM network between cells that impedes epithelial cell migration and disrupts tissue structure. These results introduce a new transport step in the BM protein secretion pathway and highlight the importance of controlling the sub-cellular exocytic site of network-forming proteins.

## RESULTS

### Khc-73 biases BM protein secretion to basal cellular regions

MTs have been implicated in polarized BM protein secretion in mammalian epithelial cells (Almeida and Stow, 1991; Boll et al., 1991), but how they influence this process is unknown. To ask if MTs play a similar role in follicle cells, we depleted them by overexpressing the MT-severing protein spastin (Sherwood et al., 2004), and visualized the BM using an endogenously GFP-tagged *α*2 chain of type IV collagen (Col IV-GFP) that produces functional Col IV (Buszczak et al., 2007). In control cells, Col IV-GFP is predominantly localized to the BM, with only small Col IV-GFP punctae along lateral surfaces (Figures 1D-F). In cells depleted of MTs, lateral Col IV-GFP is significantly increased (Figures 1D-F and S1B). Therefore, MTs are also involved in BM protein secretion in follicle cells where they bias Col IV accumulation to basal cellular regions.

We hypothesized that MTs may serve as polarized tracks for the transport of secretory vesicles filled with BM proteins toward the basal surface. Given the polarity of these MTs (Figure 1A) (Clark and Jan, 1997; Khanal et al., 2016; Nashchekin et al., 2016), this model suggests that transport should depend on a plus end-directed kinesin motor. Through an RNAi-based screen, we identified the kinesin-3 family member Khc-73 (human homolog KIF13B) as a candidate. We used CRISPR to generate a new allele, *Khc-73^3-3^*, that has an early stop codon in the motor domain (Figure S1C and Supplemental Table 3).

*Khc-73^3-3^* cells have ectopic accumulation of Col IV-GFP along lateral surfaces, similar to what we saw with spastin overexpression (Figures 1D-F). Placing the new *Khc-73^3-3^* allele in trans to the existing *Khc-73^149^* allele (Liao et al., 2018), also causes lateral Col IV-GFP accumulation (Figures S1D and S1E), confirming that mutation of Khc-73 is the source of this defect. Loss of Khc-73 similarly affects two other BM proteins, laminin and perlecan (Figure 1G). We know that the ectopic Col IV-GFP is in the extracellular space between cells because it is accessible to anti-GFP nanobodies in non-permeabilized tissue (Figure S1F). Therefore, BM proteins are secreted from *Khc-73^3-3^* cells but accumulate in the wrong location.

Our investigation of Khc-73 was motivated by the hypothesis that it transports vesicular cargo, like many kinesin-3 family members (Siddiqui and Straube, 2017). However, kinesins can play many roles in epithelial cells (Kreitzer and Myat, 2018). Apical-basal cortical polarity, the polarized localization of transmembrane proteins to apical and lateral membranes, the localization of Col IV-encoding mRNAs, and MT organization are all normal in *Khc-73^3-3^* cells (Figures 1D, S2A-D, and S3A-G). These data suggest that epithelial cell organization remains intact in the absence of Khc-73, lending support to the idea that Khc-73 affects BM protein secretion through vesicular transport.

The pattern of ectopic BM protein accumulation in *Khc-73^3-3^* epithelia is distinct from that caused by loss of previously identified regulators of polarized BM protein secretion in follicle cells, which all cause BM proteins to be mis-sorted into an apically directed secretory pathway (Denef et al., 2008; Devergne et al., 2014, 2017; Isabella and Horne-Badovinac, 2016; Lerner et al., 2013). For example, loss of Rab10 leads to abundant apical secretion of Col IV-GFP which forms a web-like network in the space between the apical surface and germ cells, with only minor Col IV-GFP accumulation along lateral surfaces (Figure 1D). In contrast, most of the ectopic Col IV-GFP in *Khc-73^3-3^* cells is below the zonula adherens (ZA), which demarcates the lateral from the apical domain (Figures 2A and 2B). The lateral Col IV-GFP accumulations appear biased to an apical (upper) region of the lateral domain below the ZA in these egg chambers, which have nearly completed new Col IV production (Figure 2A). However, if we look earlier in development when synthesis of Col IV is high, significantly more extracellular Col IV-GFP accumulates along the entire length of the lateral domain in *Khc-73^3-3^* cells within a mosaic tissue (Figures 2C-E and S1F). The biased accumulation of Col IV-GFP below the ZA later in development is likely due to the normal movement of nascent fibrils onto the BM, which clears the basal (lower) regions of the lateral domain (Figure 2F). The site of ectopic Col IV-GFP accumulation in *Khc-73^3-3^* tissue suggests that BM proteins are sorted into a basolateral trafficking pathway, but that their secretion site shifts to include upper regions of the lateral membrane (Figure 2F).

**Figure 2.**
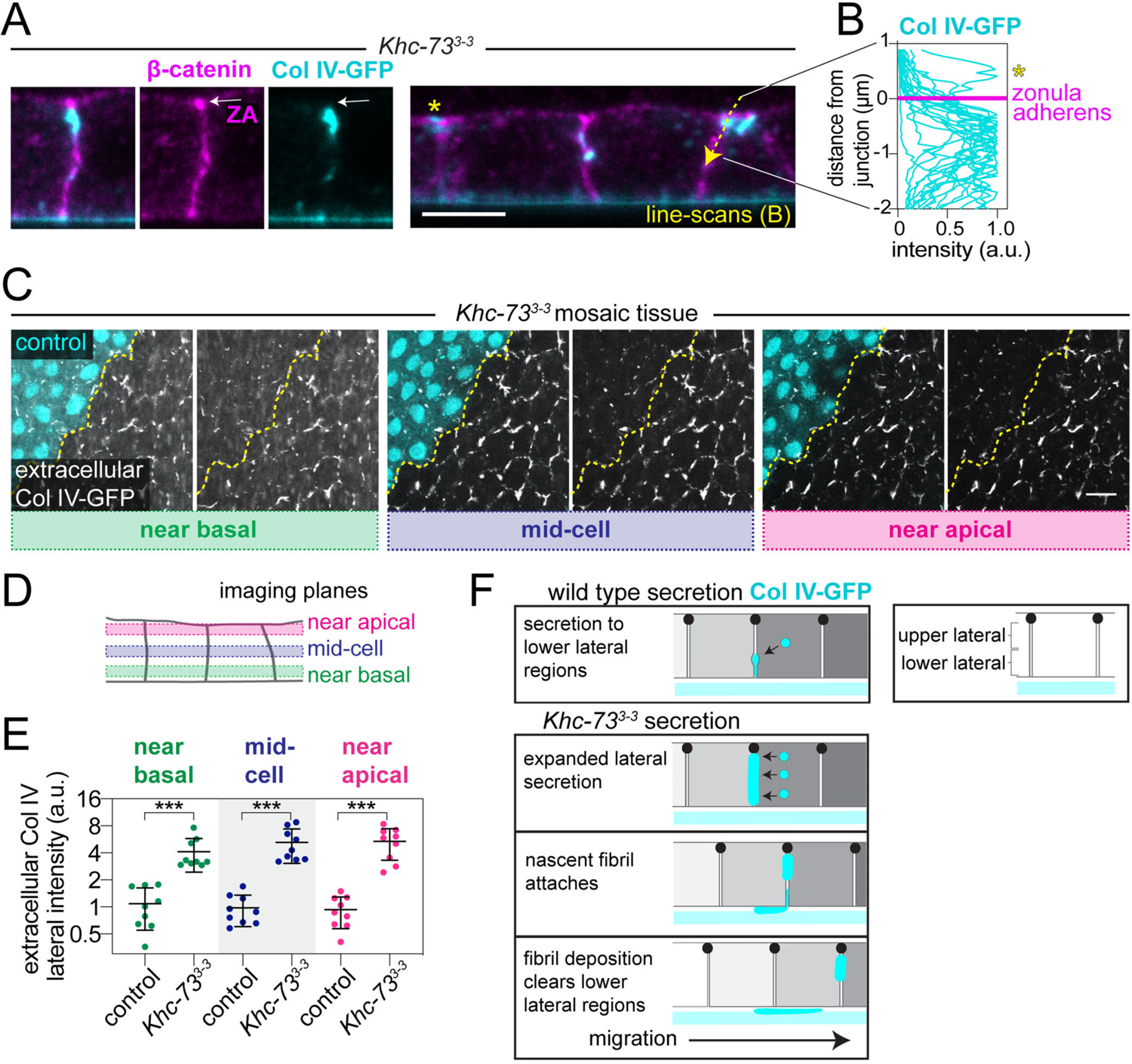
Col IV accumulates all along the lateral domain in *Khc-73^3-3^* cells. A. Images of lateral Col IV-GFP in Khc-733-3 epithelia relative to staining for the zonula adherens (ZA, white arrow). Arrow indicates where intensity line-scans were performed in (B); asterisk highlights example where Col IV-GFP is apical to the ZA. B. Graph of Col IV-GFP fluorescence intensity along 20 cell-cell interfaces as indicated in (A). Col IV-GFP line-scans were aligned to peak β-cat signal at the ZA, indicated in magenta. Asterisk highlights the example of Col IV-GFP apical to the ZA from (A). C. Images of ectopic extracellular Col IV-GFP in Khc-733-3 mosaic tissue at three different z-planes through the lateral domain, diagrammed in (D). The dotted line demarcates control and Khc-733-3 cells. Extracellular Col IV-GFP is highlighted by staining non-permeabilized tissue with a nanobody to GFP (Figure S1F). D. Illustration of imaging planes in (C). E. Quantification of extracellular Col IV-GFP in (C), showing increased lateral accumulation in Khc-733-3 cells at all three z-planes. Data represent mean ± SD plotted on a log scale. Paired t tests, ***p<0.001. n= 9 egg chambers. F. Model for how loss of Khc-73 leads to lateral Col IV accumulation, which persists mainly in the upper region of the lateral domain. Stage 8 egg chambers (A, B). Stage 7 egg chambers (C, E). Scale bars, 5 µm (A), 10 µm (B). See also Figures S1, S2, S3.

Altogether, these data suggest a model in which Khc-73 biases BM protein secretion to basal regions of the basolateral plasma membrane to facilitate their assembly into a single BM sheet at the basal surface.

### Khc-73 transports Rab10+ BM protein secretory compartments to basal regions of the cell

To determine how Khc-73 biases the site of BM protein secretion, we focused on the internal membrane compartments that mediate this process. Under normal conditions, we can only detect intracellular Col IV-GFP in the endoplasmic reticulum (ER), preventing us from directly following Col IV-GFP during secretion (Figure S4A). As an alternate marker of BM protein secretory vesicles, we used Rab10. Rabs help to define different membrane compartments by recruiting a variety of effectors (Stenmark, 2009). Rab10 directs secretory transport in other systems (Chen et al., 2012; Deng et al., 2014; Schuck et al., 2007; Zou et al., 2015), and also interacts with the human homolog of Khc-73, KIF13B (Etoh and Fukuda, 2019). In follicle cells, YFP-Rab10 localizes near the Golgi, which in *Drosophila* is distributed among all ER exit sites (ERES), and is where BM protein secretory vesicles likely form (Figures 3A and 3B) (Lerner et al., 2013). In addition, YFP-Rab10 is found on punctate and tubular compartments along the basal surface that preferentially accumulate at the trailing edge of each migrating cell (Lerner et al., 2013), where Khc-73 is also enriched (Figures 3A and 3C). As these compartments are localized away from the Golgi, they likely represent secretory intermediates.

**Figure 3.**
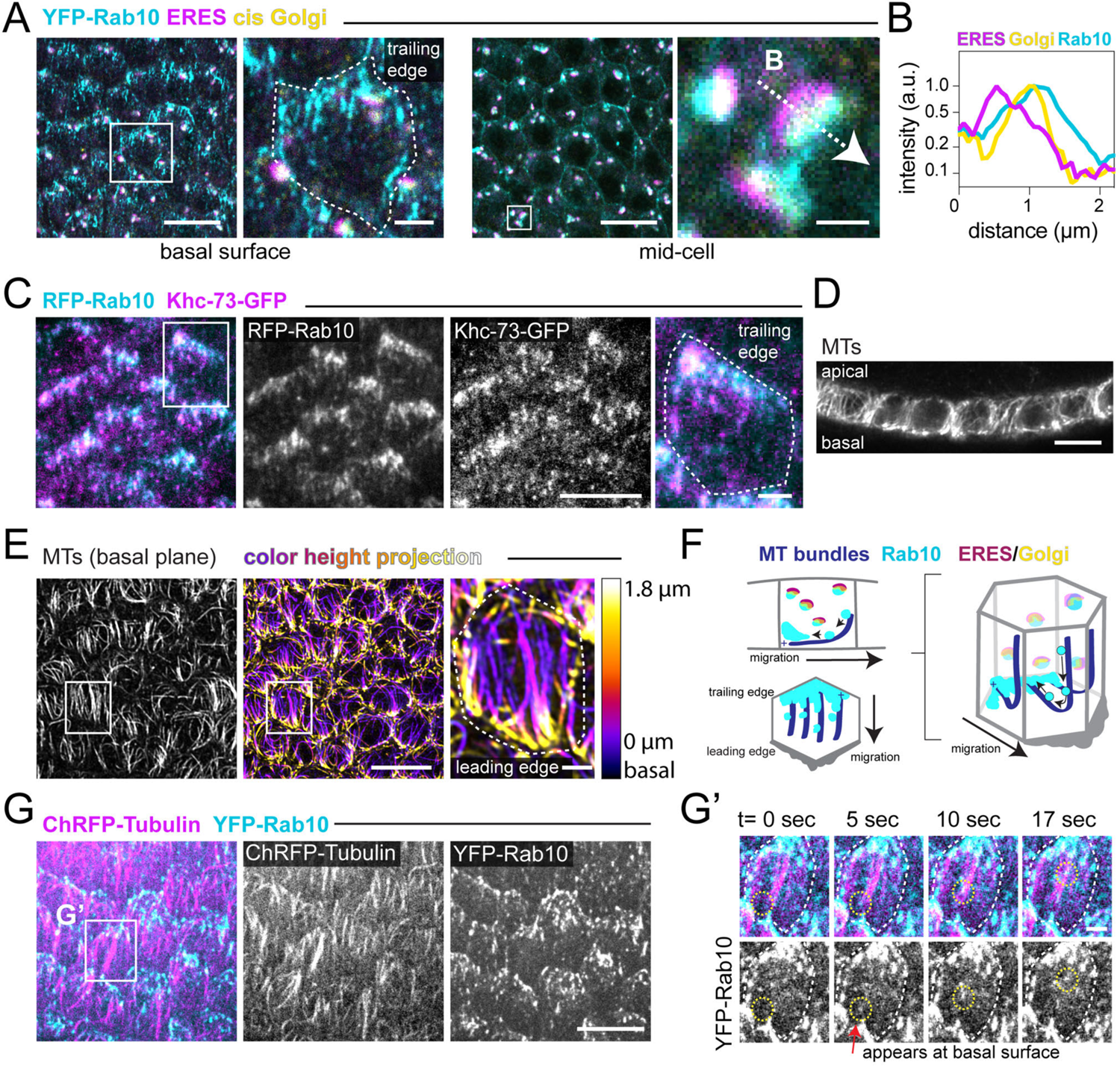
Rab10+ compartments move along MTs to basal trailing cell edges. A. Images of UAS-YFP-Rab10 localization at the basal surface and mid-cell. The mid-cell inset highlights the position of YFP-Rab10 relative to staining for the ERES protein Tango1 and the cis Golgi protein GM130 (line-scan of example intensity profiles in B). The basal inset highlights one cell traced with a dotted line with UAS-YFP-Rab10 at trailing edge. Scale bars, 10 µm main panels, 2 µm basal inset, and 1 µm ERES/Golgi inset. B. Line-scan of fluorescence intensity along the arrow in (A). C. Image of Khc-73-GFP (endogenous promoter) localizing to UAS-RFP-Rab10+ tubular compartments at the basal trailing edges of cells. Inset cell traced with dotted line. D. Image showing MTs (anti-acetylated ⍺-tubulin) aligned along the apical-basal axis in a cross-section through follicle cells. E. Image of MTs (anti-acetylated ⍺-tubulin) aligned parallel to the direction of migration along the basal surface (left panel). A color height projection (center panel) of the basal-most 1.8 μm of the epithelium shows that the basal MT bundles at the leading edges of cells bend and extend apically. Inset highlights a single cell traced with a dotted line. See also Movie 2. F. Illustration of the 3D organization of Rab10+ compartments and MT bundles that bend near the basal leading cell edges, as viewed in: cross-section, along the basal surface, or in a 3D cell. Only a few examples of this bent population of MTs/MT bundles are shown for clarity. We could not follow all MTs in 3D in all areas of cell. G. Image from time-lapse showing that UAS-YFP-Rab10+ compartments colocalize with and move along the basal MT array (UAS-ChRFP-⍺-tubulin). (G’) Montage of YFP-Rab10+ puncta appearing in the focal plane and moving along a MT bundle from inset in G. See also Movie 3. Stage 7 egg chambers (A, B, C, D, E). Stage 8 egg chamber (G). Images oriented such that migration is down. Scale bars, 10 µm for main panels and 2 µm for cell insets, except as described in (A). See also Figure S4A, Movie 2, and Movie 3.

To understand the organization of the MTs that may be used to transport Rab10+ compartments toward the basal surface, we examined them in 3D. Similar to other epithelia, one MT array runs parallel to the apical-basal axis of follicle cells (Figure 3D), with minus ends anchored apically and plus ends growing toward the basal surface (Figure 1A) (Clark and Jan, 1997; Khanal et al., 2016; Nashchekin et al., 2016). Follicle cells also have a MT array along their basal surfaces that is thought to be involved in collective migration (Chen et al., 2016; Viktorinová and Dahmann, 2013). These MTs are aligned parallel to the direction of migration (Figure 1E), and their plus ends grow preferentially toward trailing cell edges (Figures 1A) (Viktorinová and Dahmann, 2013). In 3D image volumes taken near the basal surface, we found a connection between these two MT arrays. The basal MT bundles lie flat along part of the basal surface, but bend sharply near the leading edge of cells, integrating seamlessly into the apical-basal array (Figure 3E and Movie 2). This MT organization could allow kinesins to bring Rab10+ secretory vesicles from all over the cell to the basal trailing edge where Rab10+ compartments are enriched (Figure 3F).

To determine if Rab10+ punctae move directionally along MTs as expected for a kinesin cargo, we focused on the basal surface because its location on the exterior of the tissue provides superior imaging quality and we can follow the population of Rab10+ punctae that likely mediate basally polarized secretion. YFP-Rab10+ punctae localize to MTs and move rapidly along them (Figure 3G and Movie 3). We frequently observed motile punctae enter the focal plane at leading edges in regions where a MT bundle also came into view (Figure 3G’ and Movie 2), suggesting that these bent MTs are used for transport from more apical regions to the basal surface. Finally, we tracked the direction of rapidly moving YFP-Rab10+ punctae and found that they moved toward the trailing edge of cells 2.12-fold more often than toward the leading edge (297 trajectories in 5 egg chambers, data also used as control in Figures 4F-H). This directional bias is consistent with a role for MT plus end-directed transport of Rab10+ secretory vesicles to basal trailing cell edges (Figure 3F).

**Figure 4.**
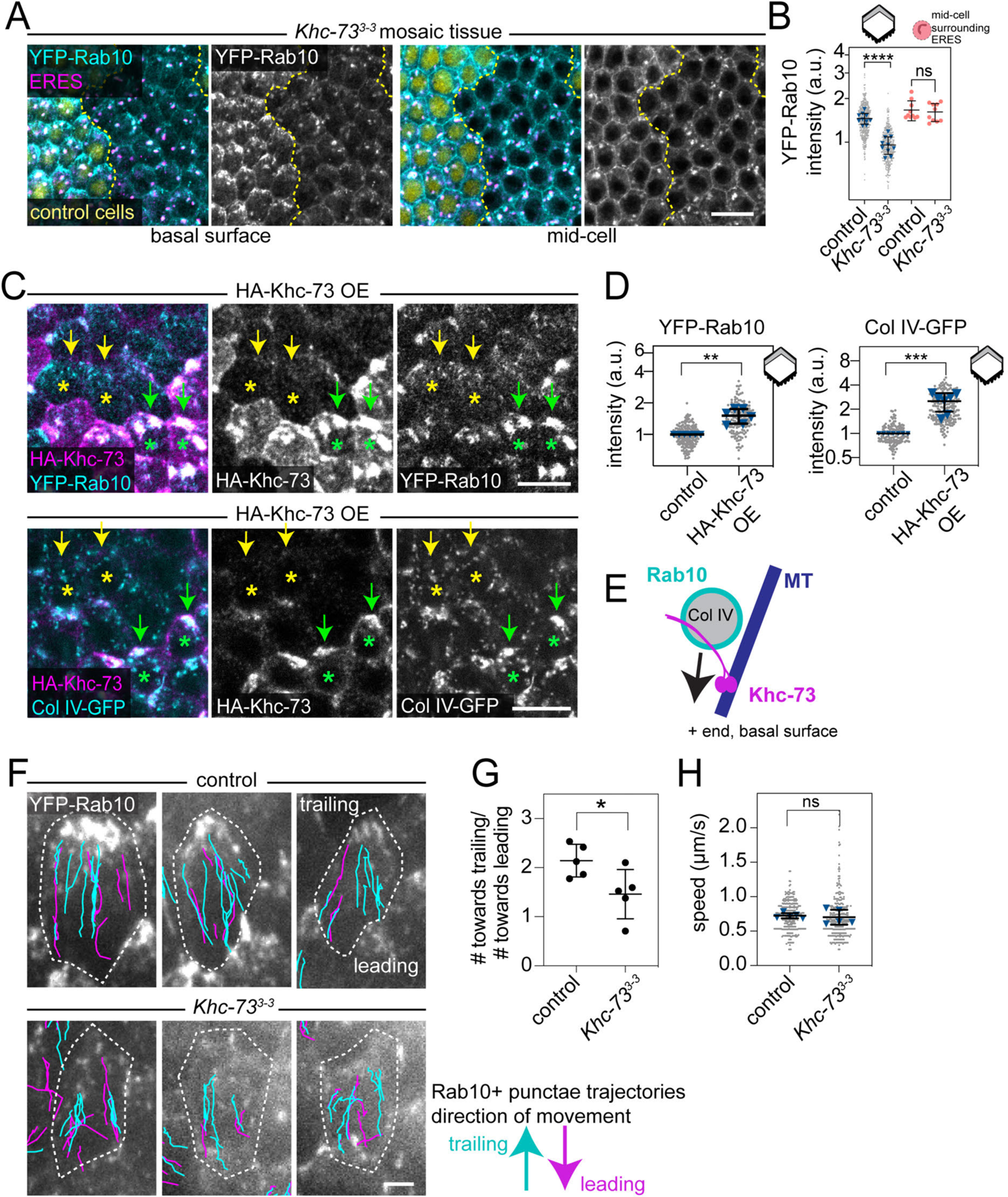
Khc-73 transports Rab10+ compartments to the basal trailing edges of follicle cells. A. Images of UAS-YFP-Rab10 and ERES (anti-Tango1) in *Khc-73^3-3^* mosaic tissue at the basal surface and mid-cell. The dotted line demarcates control and *Khc-73^3-3^* cells. B. Quantification of the decrease in UAS-YFP-Rab10 levels at basal trailing cell edges (grey region of cell in cartoon) without a change in UAS-YFP-Rab10 levels near the ERES mid-cell (salmon) in *Khc-73^3-3^* cells. Grey dots represent individual cells and blue triangles represent egg chamber means. For basal surface: paired t test, ****p<0.0001. n=10 egg chambers, 544 control and 457 *Khc-73^3-3^* cells. For ERES: Wilcoxon matched-pairs signed rank test, ns p>0.05. n=10 egg chambers. C. Images of YFP-Rab10 (endogenous, top panels), and Col IV-GFP (endogenous, bottom panels) in epithelia overexpressing UAS-HA-Khc-73 in patches of cells. YFP-Rab10 images are along basal surface, and Col IV-GFP panels are 1.5 µm above the basal surface to avoid the Col IV-GFP signal within the BM. Yellow asterisks indicate “control” cells negative for HA-Khc-73 staining; yellow arrows point at the trailing cell edges. Green asterisks label HA-Khc-73 OE cells; green arrows point at the punctae accumulating at trailing cell edges. See also Movie 4. D. Quantification of the increase in YFP-Rab10 (endogenous) and Col IV-GFP (endogenous) in HA-Khc-73 OE cells at basal trailing cell edges (grey region of cell in cartoon). Grey dots represent individual cells and blue triangles represent egg chamber means. One sample t tests compared to the theoretical ratio of 1, **p<0.01, ***p<0.001. For YFP-Rab10, n=6 egg chambers, 199 “control” and 161 HA-Khc-73 OE cells. For Col IV-GFP, n=7 egg chambers, 149 “control” and 214 HA-Khc-73 OE cells. E. Model of Khc-73’s role in transporting Rab10+ Col IV-filled secretory vesicles toward the basal surface. F. Images of example cells (dashed outlines) from the first frame of a time-lapse of control and *Khc-73^3-3^* epithelia expressing UAS-YFP-Rab10. Trajectories of UAS-YFP-Rab10+ punctae are overlaid on the images and color-coded for direction. See also Movie 5. G. Quantification of the direction of UAS-YFP-Rab10+ punctae movements from (F) scored as either toward the trailing edge or the leading edge in control and *Khc-73^3-3^* epithelia. Unpaired t test, *p<0.05. n= 5 egg chambers, 297 runs in control and 305 runs in *Khc-73^3-3^* egg chambers. H. Distribution of speeds of UAS-YFP-Rab10+ punctae in control and *Khc73^3-3^* epithelia at the basal surface. Grey dots represent individual runs and blue triangles represent egg chamber means. Unpaired t test, ns p>0.05. n= 5 egg chambers, 297 runs in control and 305 runs in *Khc-73^3-3^* egg chambers. Stage 7 egg chambers. Images oriented such that migration is down. Data represent mean ± SD. Statistics were performed on egg chamber mean values. Data in B and D are plotted on a log scale. Scale bars, 10 µm (A and C), 2 µm (F). See also Figure S4B, S4C, Movie 4, and Movie 5.

We next asked if Khc-73 is the motor that transports Rab10+ compartments basally. Loss of Khc-73 reduces YFP-Rab10 intensity at basal trailing cell edges, but not at the Golgi (Figures 4A and 4B). This result is consistent with our observations of extracellular Col IV-GFP accumulation that suggest Khc-73 is not needed for secretion to occur, just to specify the correct location (Figures 2C-E). Conversely, overexpressing Khc-73 increases YFP-Rab10 intensity at basal trailing cell edges, where it forms large, aberrant foci (Figures 4C and 4D). We do not normally see Col IV-GFP colocalized with Rab10 in control cells. However, Col IV-GFP is concentrated within the aberrant foci induced by Khc-73 overexpression (Figures 4C and 4D), which suggests the foci are clusters of trapped BM protein vesicles. In support of this idea, the foci lack markers of earlier secretory compartments like the ER and Golgi (Figures S4B and S4C), and YFP-Rab10+ tubulovesicular structures move rapidly into and out of the foci (Movie 4). These data show Khc-73 is necessary for Rab10+ compartment localization to the basal surface, and sufficient to alter the localization of Rab10+ and Col IV+ compartments when over-expressed.

Altogether, our observations that Rab10+ punctae move preferentially toward cellular regions enriched in growing MT plus-ends, and that changes in Khc-73 expression affect the localization of both Rab10+ and Col IV+ compartments, strongly suggest that Khc-73 transports BM protein secretory vesicles basally to their secretion site (Figure 4E).

### Kinesin-1 works with Khc-73 to direct BM protein secretion basally

Khc-73 is the only motor whose individual knock-down perturbed Col IV secretion in our RNAi screen. However, live imaging and tracking of YFP-Rab10+ punctae movements in *Khc-73^3-3^* tissue revealed that the YFP-Rab10+ punctae that do reach the basal surface move at normal speeds (control: 0.72 ± 0.21 and *Khc-73^3-3^*: 0.71 ± 0.31 µm/s), although with slightly less directional bias toward the trailing edge of cells (Figures 4F-H, and Movie 5). When multiple kinesins contribute to transport, the cargo speed can be dominated by one motor (Arpağ et al., 2014, 2019; Norris et al., 2014). Since kinesin-1 transports secretory vesicles to specify the secretion site of many proteins (Fourriere et al., 2019), and more specifically transports Rab10+ vesicles in mammalian neurons (Deng et al., 2014), we investigated if *Drosophila’s* sole kinesin-1 motor, Khc, might work with Khc-73 to transport Rab10+ BM protein secretory vesicles.

To test this hypothesis, we depleted both kinesins. Compared to depletion of Khc-73 alone, co-depletion of Khc-73 and Khc using RNAi caused Col IV-GFP to accumulate at lateral cell edges at higher levels (Figures 5A-C). We confirmed that loss of Khc alone does not cause lateral Col IV-GFP accumulation using a Khc null mutation, *Khc^27^* (Figures S5A and S5B) (Brendza et al., 1999). Co-depletion of Khc-73 and kinesin light chain (Klc), which often acts as a cargo adaptor for Khc (Kamal and Goldstein, 2002), similarly enhanced lateral Col IV-GFP accumulation (Figures 5A-C). These dual RNAi experiments uncovered that Khc and Klc work with Khc-73 to promote basally polarized BM protein secretion.

**Figure 5.**
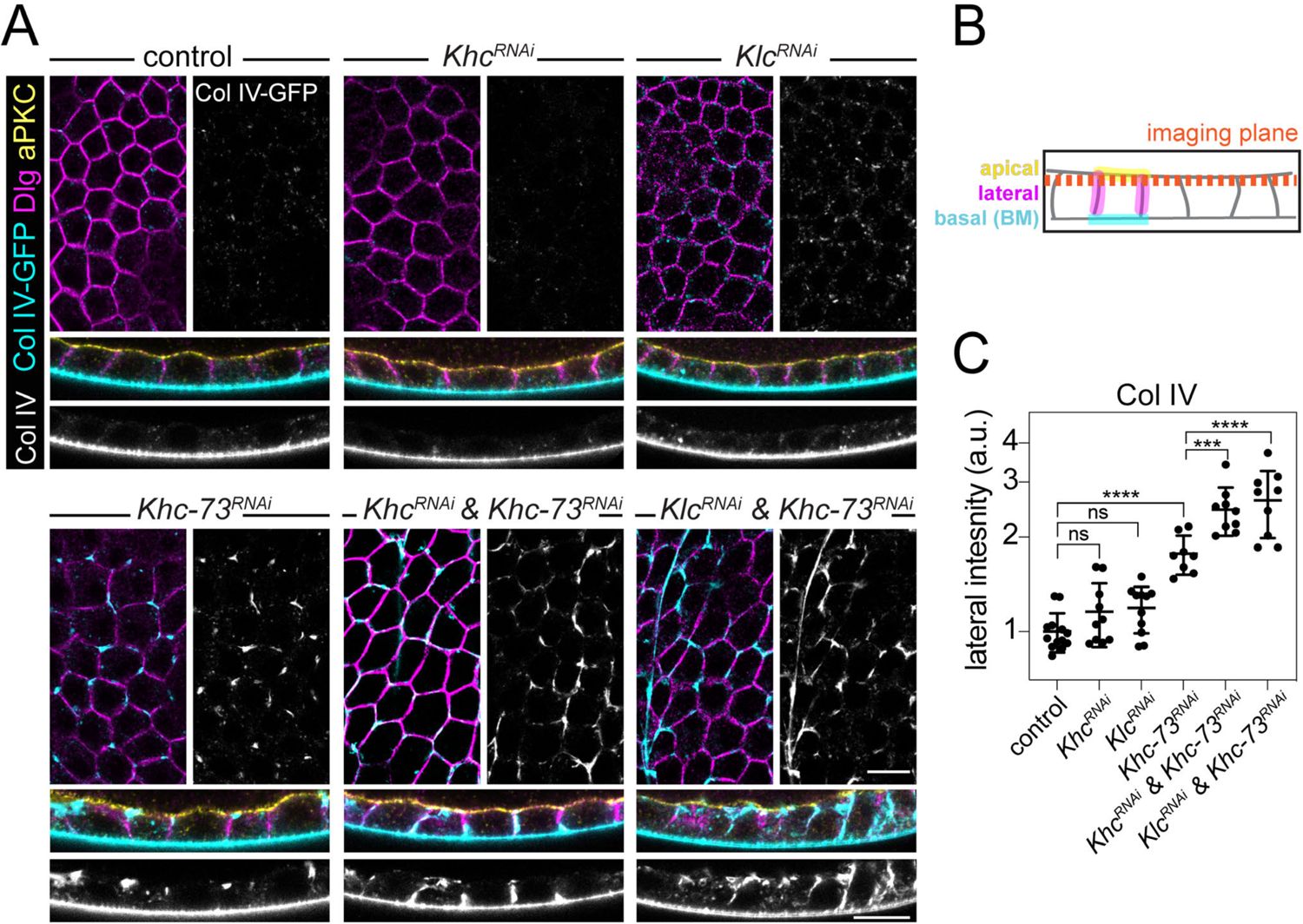
The kinesin-1 Khc works with Khc-73 to direct polarized BM protein secretion. A. Images showing ectopic Col IV-GFP accumulation in epithelia expressing RNAi against Khc, Klc, Khc-73, and combinations thereof. Top panels are planes through the lateral domains that capture some of the apical surface due to tissue curvature, as diagrammed in (B). Bottom panels are cross sections. Anti-Dlg marks lateral domains. Anti-aPKC marks apical domains (only shown in cross sections). B. Illustration of imaging planes in (A). C. Quantification of ectopic Col IV-GFP accumulation at lateral cell edges in the plane used in (A) for all RNAi conditions. Data represent mean ± SD plotted on a log scale. Ordinary one-way ANOVA with Holm-Šídák’s multiple comparisons test; ns p>0.05, ***p<0.0005, ****p<0.0001. In the order on graph, n=13, 10, 11, 8, 9, 9 egg chambers. Stage 8 egg chambers. Scale bars, 10 µm. See also Figure S5A and S5B.

We next asked if Khc affects BM protein secretion by the same mechanism as Khc-73, transporting Rab10+ compartments toward the basal surface. Since Khc loss alone is insufficient to disrupt the site of polarized Col IV-GFP secretion (Figures S5A and S5B), we expected a milder reduction in YFP-Rab10+ compartments at the basal trailing edges of *Khc^27^* cells than in *Khc-73^3-3^* cells (Figures 4A and 4B). Surprisingly, this population of YFP-Rab10+ compartments is instead increased in *Khc^27^* cells (Figures 6A and 6B). Khc has two roles that could impact the localization of Rab10+ compartments; Khc transports vesicles and also rearranges the MT tracks themselves by binding a second MT with its tail domain and sliding the two MTs relative to each other (Jolly et al., 2010). MTs remain aligned in *Khc^27^* cells (Figures S5C-E), and a mutation in Khc’s tail domain that selectively impairs its ability to slide MTs does not affect YFP-Rab10+ compartment localization (Figures 6A and 6B) (Winding et al., 2016). These data point to Khc’s cargo transport activity contributing to the normal localization pattern of Rab10+ compartments.

**Figure 6.**
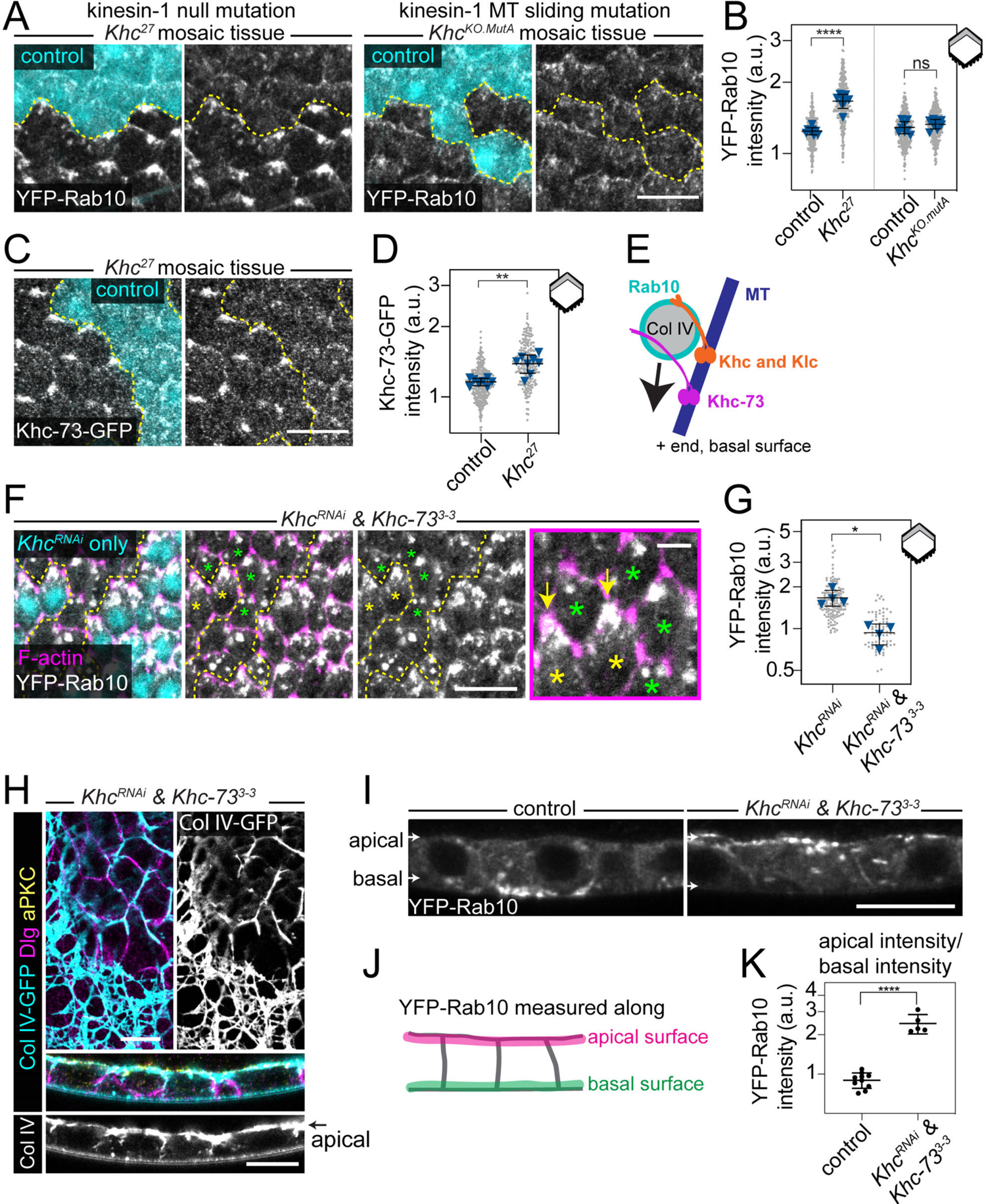
Khc contributes to the transport of Rab10+ compartments to the basal surface. A. Image of YFP-Rab10 (endogenous) along the basal surface in Khc27 and KhcKO.mutA mosaic tissues. The dotted line demarcates control and mutant cells. B. Quantification of the increase in YFP-Rab10 (endogenous) at the basal trailing edge (grey region of cell in cartoon) of Khc27 cells, with no change in KhcKO.mutA cells. Grey dots represent individual cells and blue triangles represent egg chamber means. Paired t tests, ns p>0.05, ****p<0.0001. For Khc27, n=8 egg chambers, 351 control and 339 mutant cells. For KhcKO.mutA, n=8 egg chambers, 344 control and 366 mutant cells. C. Image of Khc-73-GFP (endogenous promoter) along the basal surface in Khc27 mosaic tissue. The dotted line demarcates control and Khc27 cells. D. Quantification of the increase in Khc-73-GFP (endogenous promoter) at the basal trailing edge (grey region of cell in cartoon) of Khc27 cells. Grey dots represent individual cells and blue triangles represent egg chamber means. Paired t test, **p<0.01. n=8 egg chambers, 372 control and 245 Khc27 cells. E. Model for two kinesins’ role in transporting BM protein secretory vesicles. F. Image of UAS-YFP-Rab10 along the basal surface in KhcRNAi & Khc-733-3 mosaic tissue. The dotted line demarcates the KhcRNAi only and KhcRNAi & Khc-733-3 cells. F-actin is shown to visualize cell edges. Yellow asterisks mark two KhcRNAi only cells and green asterisks mark the KhcRNAi & Khc-733-3 cells enlarged in inset (magenta outline) to highlight higher YFP-Rab10 accumulation at trailing edges (arrows) of KhcRNAi only cells. G. Quantification of the decrease in UAS-YFP-Rab10 in KhcRNAi & Khc-733-3 cells relative to KhcRNAi only cells at basal trailing edges (grey region of cell in cartoon). Grey dots represent individual cells and blue triangles represent egg chamber means. Paired t test, *p<0.05. n=4 egg chambers, 153 KhcRNAi only and 79 KhcRNAi & Khc-733-3 cells. H. Images of the ectopic apical Col IV-GFP network formed in a KhcRNAi & Khc-733-3 epithelium. Top panel is a plane through the lateral domains that captures some of the apical surface due to tissue curvature. Bottom panel is a cross section. Anti-Dlg marks lateral domains. Anti-aPKC marks apical domains (only shown in cross sections). Same imaging and display settings as Figure 5A. I. Images of UAS-YFP-Rab10 in cross-sections through control and KhcRNAi & Khc-733-3 epithelia. In control, UAS-YFP-Rab10 at the basal surface is only visible where the cross-section passes though the trailing edge of a cell. Apical UAS-YFP-Rab10 is uniformly along the apical surface in mutant. J. Illustration of where YFP-Rab10 intensity was measured in (I). K. Quantification of the apical surface enrichment of UAS-YFP-Rab10 in KhcRNAi & Khc-733-3 epithelia, measured along the surfaces illustrated in (J). Data represent mean ± SD and are plotted on a log scale. Unpaired t test, ****p<0.0001. n=10 control and 5 KhcRNAi & Khc-733-3 egg chambers. Stage 7 egg chambers (A-G). Stage 8 egg chambers (H-K). Data represent mean ± SD and are plotted on a log scale (B,D,G). Statistics are performed on egg chamber means. Scale bars 10 µm, except inset in F which is 2 µm. See also Figures S5C-E and S6.

The increase in Rab10+ compartments at the basal surface in *Khc^27^* mutant cells was unexpected. We hypothesized that loss of Khc is compensated for by increased Khc-73 activity, and that high Khc-73 activity changes Rab10+ compartment localization. In support of this idea, the increase in YFP-Rab10+ compartments at the basal trailing edges of *Khc^27^* cells resembles the aberrant YFP-Rab10+ foci that form when Khc-73 is overexpressed (Figures 4C and 4D). In addition, Khc-73-GFP levels are increased at the basal trailing edges of *Khc^27^* cells (Figures 6C and 6D). Finally, removal of both kinesins prevented nearly all YFP-Rab10+ compartment accumulation at basal trailing edges (Figures 6F and 6G), showing that Khc-73 mediates the increase. These data suggest that while Khc normally contributes to the transport of Rab10+ BM protein secretory vesicles (Figure 6E), Khc-73 can largely compensate for its loss. However, increased reliance on Khc-73 leads to more Rab10+ compartment localization to basal trailing cell edges (Figures 6A and 6B), revealing a distinct role for Khc in achieving the normal distribution of Rab10+ compartments.

Further reducing kinesin levels by expressing *Khc^RNAi^* in *Khc-73^3-3^* cells, as we did to visualize Rab10 above, produced a new phenotype; Col IV-GFP accumulated on the apical surface (Figure 6H). In these cells, YFP-Rab10+ compartments are not only lost from the basal surface but also become concentrated beneath the apical surface (Figures 6I-K). Importantly, general cell organization is unaffected in *Khc^RNA^*^i^ & *Khc-73^3-3^* cells, as the polarized localization of cortical and transmembrane proteins, Col IV-encoding mRNAs and MTs all remain normal (Figures 6H and S6A-G). Therefore, these two kinesins not only contribute to post-Golgi BM protein secretory vesicle transport, but also play an essential role in preventing apical secretion of BM proteins that is only revealed when they are severely depleted.

### Intercellular BM protein networks disrupt epithelial architecture

Finally, we investigated how changing the subcellular secretion site of BM proteins impacts the final structure of the BM and the follicle cells migrating along it. We initially focused on the BM sheet along the basal surface. The mean level of Col IV-GFP in the BM is normal in all genotypes where kinesins are reduced, except the condition that causes apical BM protein secretion, *Khc-73^3-3^ & Khc^RNAi^* (Figures 7A and B). However, there are striking changes to the structure of the BMs in all conditions. There are more and larger fibrils in *Khc-73^3-3^* BMs and *Khc-73^RNAi^ & Khc^RNAi^* BMs (Figures 7A and 7D). Therefore, in these kinesin mutant conditions, although Col IV is initially mis-secreted all along the lateral membrane, much of this protein is eventually moved onto the underlying BM as fibrils (Figure 2F).

**Figure 7.**
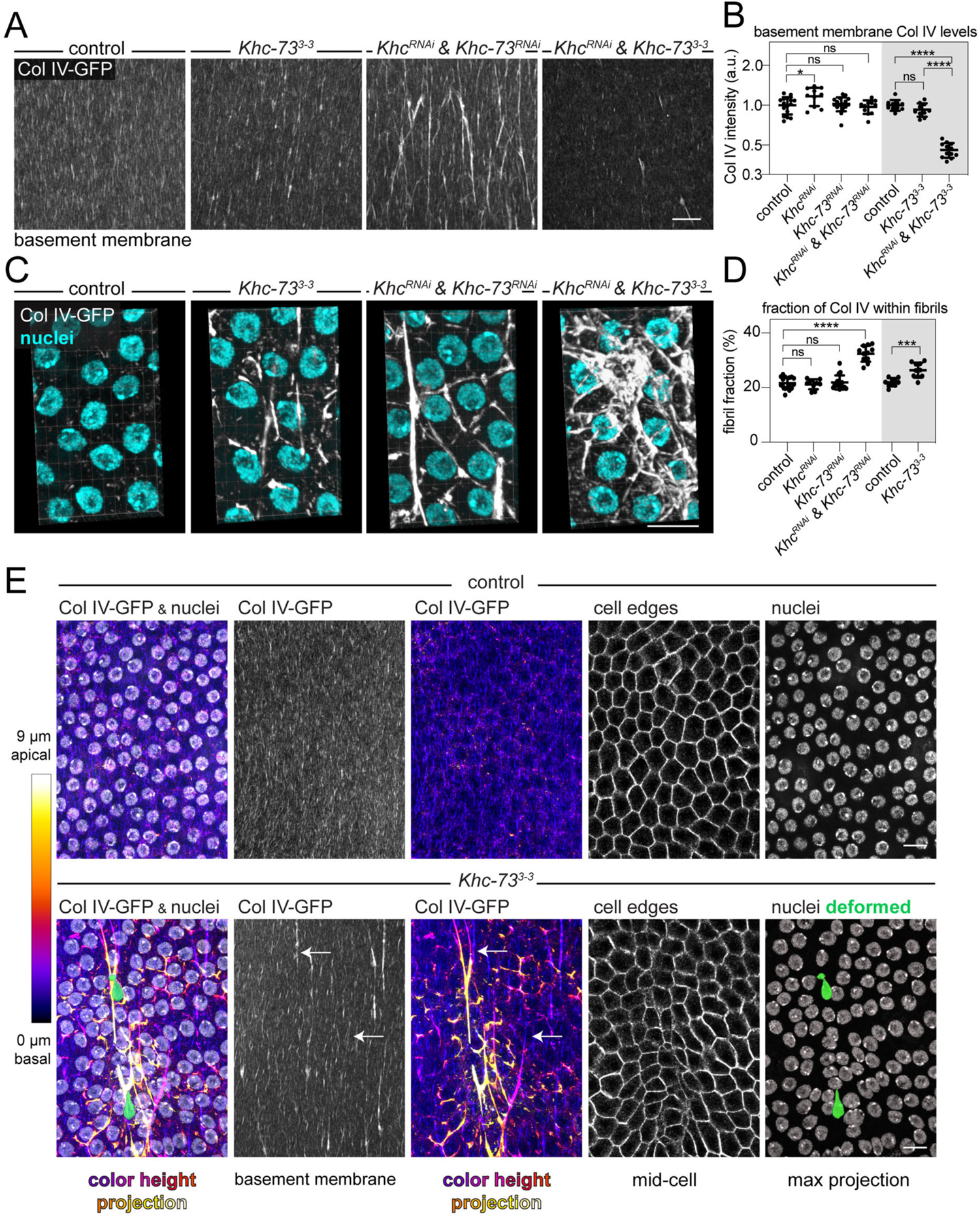
Intercellular BM protein networks disrupt epithelial architecture. A. Images showing changes in the organization, and intensity, of Col IV-GFP within the BMs of kinesin mutant egg chambers. B. Quantification of mean Col IV-GFP intensity in the BMs from (A). Ordinary one-way ANOVA with Šídák’s multiple comparisons test; ns p>0.05, *p<0.05, ****p<0.0001. In order on graph, n=16,10,16,11,10,10,12 egg chambers. C. 3D projection of Col IV-GFP intercellular networks formed in a Khc-733-3 epithelium and a Khc-73RNAi & KhcRNAi epithelium. An apical web-like network is formed in a KhcRNAi & Khc-733-3 epithelium. The planes containing the basal sheet of BM were removed before making projection. The view shown is looking down onto the apical surfaces of follicle cells. D. Quantification of the fraction of Col IV-GFP intensity associated with BM fibrils in (A). One-way ANOVA followed by Dunn’s multiple comparison test, ns p>0.05, ****p<0.0001. For control and Khc-733-3 (grey region of graph), unpaired t test, ***p<0.0005. In order on graph, n=16,9,16,11,10,10 egg chambers. E. Images showing the effect of Col IV-GFP intercellular cables on cell and nuclear shapes. A color height projection of Col IV-GFP from a confocal volume of the full-thickness of the follicle cells shows ectopic cables of Col IV-GFP pass through the region where cells and nuclei are deformed. Arrows indicate where intercellular Col IV cables contact the underlying BM. Images oriented such that migration is down. See also Movie 6 Stage 8 egg chambers. Data represent mean ± SD and are plotted on a log scale. Scale bars 10 µm.

We know that these kinesin mutant conditions also have persistent lateral accumulations of Col IV. To understand how this Col IV is organized relative to follicle cells, we made 3D reconstructions of the Col IV-GFP that is not in the BM. In reconstructions of the apical secretion condition, *Khc-73^3-3^ & Khc^RNAi^*, a web-like Col IV-GFP network lies atop the apical surface of follicle cells (Figure 7C). In contrast, in conditions that cause lateral secretion, Col IV-GFP forms an intercellular network that spans multiple cell lengths (Figure 7C). Some regions of these lateral, intercellular networks are connected to the underlying BM. As follicle cells are collectively migrating relative to the BM, these connections likely act as anchor points that impede cell movement. This is best illustrated in the mildest condition, *Khc-73^3-3^*, where there are only a few, isolated Col IV-GFP cables running between cells and anchored to the BM at one end (Figure 7E, arrows, and Movie 6). The hexagonal packing of follicle cells is disrupted specifically near these cables compared to other areas of the epithelium (Figure 7E). Even more strikingly, several nuclei are highly deformed (Figure 7E). Since follicle cells collectively migrate in a tightly packed sheet, some cells are likely forced to squeeze through the intercellular Col IV-GFP network, causing this nuclear deformation. This drastic effect on cells from only a relatively small fraction of Col IV within these cables highlights the importance of tightly controlling the secretion site of these network-forming proteins.

## DISCUSSION

This work provides the first mechanistic insight into how the secretion site of BM proteins is controlled in an epithelium. Like mammalian epithelia, we found that MTs influence polarized BM protein secretion in follicle cells. We further identified a kinesin-3 motor, Khc-73, and a kinesin-1 motor, Khc, required for this process. When these motors are depleted, BM proteins form ectopic networks between follicle cells. Reducing kinesin levels also alters the localization and movement of Rab10+ BM protein-filled secretory compartments. The impact of kinesin depletion on the localization of both intracellular Rab10+ compartments, and extracellular ectopic Col IV accumulations, lead us to propose that these two kinesins transport BM protein-filled secretory vesicles towards the basal surface to bias the site of BM protein secretion to basal cellular regions.

How are kinesins recruited to BM protein secretory vesicles? We previously showed that Rab10 is required to sort BM proteins into a basally directed trafficking pathway and that it localizes near the Golgi where secretory vesicles likely form (Lerner et al., 2013). Based on Rab10’s role in trafficking other types of secretory vesicles (Chen et al., 2012; Deng et al., 2014; Schuck et al., 2007; Zou et al., 2015), it is likely that Rab10 remains associated with BM protein secretory vesicles during transport to the cell surface, recruiting different effectors over time to execute subsequent trafficking steps. We envision that Rab10 recruits the kinesins to newly formed BM protein-filled vesicles near the Golgi to initiate their basally-directed transport. In mammalian cells, Rab10 interacts with both kinesins we identified. Rab10 directly binds to Khc-73’s homolog, KIF13B (Etoh and Fukuda, 2019), and indirectly associates with kinesin-1 via the adaptor protein Jip1 and kinesin light chain (Deng et al., 2014). Rab8 is also required for polarized BM protein secretion in follicle cells (Devergne et al., 2017), which raises the possibility that Rab8 recruits the kinesins either alone or in combination with Rab10. To date, however, Rab8 has not been linked to either kinesin in this study. Since kinesins need to not only be recruited, but activated by association with their cargo or other proteins such as adaptors (Fu and Holzbaur, 2014), distinguishing between conservation of these mammalian interactions and other models for how the kinesins are recruited to BM protein vesicles will be an important area for future research.

The involvement of both kinesin-3 and kinesin-1 family motors in the transport of Rab10+ compartments raises questions about their respective contributions. Individual loss of either Khc or Khc-73 has the opposite effect on Rab10+ compartment localization to basal trailing cell edges, suggesting the two motors make distinct contributions to transport of Rab10+ compartments. Studies of how other kinesin-3 and kinesin-1 family motors contribute to transport when on the same cargo provide a useful framework to think about how they may work together in follicle cells. Kinesins are placed under load during transport by pulling against their cargo, and each other. The ability of kinesin-1 motors to remain bound to MTs better under load than kinesin-3 motors allows kinesin-1 to play a dominant role in the movement of cargo (Arpağ et al., 2014, 2019; Norris et al., 2014). Kinesin-3 family motors detach from MTs easily under load, but they also rebind very quickly, leading to the model that kinesin-3 motors allow cargo to remain attached to MTs in the face of obstacles in the cell (Arpağ et al., 2019; Budaitis et al., 2021; Norris et al., 2014). This, along with kinesin-3’s higher speeds and longer run-lengths, allows kinesin-3 motors to facilitate long distance transport (Huckaba et al., 2011; Siddiqui and Straube, 2017; Soppina et al., 2014). These differences in kinesin-1 and kinesin-3 behavior are one important determinant of their contributions to collectively transporting cargo.

In follicle cells, loss of Khc-73 alone disrupts polarized BM protein secretion, while loss of Khc alone does not, suggesting the kinesin-3, Khc-73, plays the major role in secretory transport. However, when Khc-73 is removed, although there are fewer Rab10+ punctae at the basal surface, they move at normal speeds. If Khc-73 and Khc are both on the same Rab10+ vesicle, Khc’s superior ability to remain attached to MTs under load may allow it to play a dominant role in cargo movement even under normal conditions, and it simply continues to transport Rab10+ vesicles when Khc-73 is missing. Why then are Rab10+ compartments reduced at the basal surface when Khc-73 is missing? Since kinesin-3 family motors facilitate long distance transport, Khc-73 loss may reduce the ability of Rab10+ compartments to initially associate with, or remain attached to, MTs. Conversely, when Khc-73 operates without Khc, it not only compensates for Khc’s loss but transports more Rab10+ compartments to basal trailing cell edges than normal. Since this location is where growing MT plus ends terminate (Viktorinová and Dahmann, 2013), this is potentially also explained by Khc-73 mediating longer-distance movements of Rab10+ compartments than Khc. Since we only image the basal surface of each cell, we rarely observed the beginning and end of a Rab10+ puncta trajectory, precluding analysis of run lengths. Following the dynamics of Rab10+ compartments in 3D would allow more comprehensive analysis of their movements, and shed new light on how the two kinesins mediate Rab10+ transport from all over the cell to the basal trailing cell edge.

A recent study on the human homologs of Khc-73 and Khc, KIF13B and KIF5B, thoroughly dissected their distinct contributions to the transport of Rab6+ secretory vesicles in non-polarized HeLa cells (Serra-Marques et al., 2020). The authors found that both kinesins are needed to bring secretory vesicles all the way to the plus ends of dynamic MTs, spatially determining where secretion occurs (Serra-Marques et al., 2020). This is similar to how we think kinesins specify the site of BM protein secretion. However, in HeLa cells the more highly expressed kinesin-1 motor plays the major role in determining where Rab6+ vesicles fuse with the plasma membrane (Serra-Marques et al., 2020). This difference highlights the importance of studying the transport of a variety of native cargos *in vivo,* where differences in motor expression, cell polarity, and other key regulators of kinesins like MT-associated proteins and MT modification can influence transport (Guardia et al., 2016; Guedes-Dias et al., 2019; Monroy et al., 2020; Norris et al., 2014; Reed et al., 2006).

Our work also introduces an unusual architecture for an epithelial MT array. Previous studies described two polarized MT arrays in the follicle cells – one running from the apical surface to the basal surface, which is standard in epithelia (Clark and Jan, 1997; Khanal et al., 2016; Nashchekin et al., 2016), and another along the basal surface of the cells, which has been linked to regulation of collective migration (Chen et al., 2016; Viktorinová and Dahmann, 2013). We find these two MT arrays are interconnected, which suggests that MT motors may move cargo seamlessly between the apical-basal and planar MT arrays. Because MT plus ends grow both toward the basal surface and toward the trailing edges of cells, this connection could explain why Rab10+ compartments are enriched at basal trailing edges.

Whether this MT arrangement leads to planar-polarized secretion of BM proteins has not yet been directly demonstrated, but we envision that such a mechanism would bring BM protein secretory vesicles all the way to the trailing edge of cells where they may naturally fuse with the lower lateral membrane for secretion to promote BM fibril formation. Although MT motors are known to be important for the secretion of many apical proteins in epithelial cells, little is known about the role of MTs and MT motors in the secretion of basolateral proteins (Rodriguez-Boulan and Macara, 2014). Determining how this MT array is built, and the significance that it holds for polarized membrane traffic, is likely to reveal the strategies cells employ to specify discrete secretion sites within larger plasma membrane domains. In the migrating follicle cells, this will also further our understanding of how cells coordinate migration with secretion.

Our finding that a moderate depletion of kinesins causes BM proteins to accumulate between the lateral membranes of follicle cells is significant because this phenotype is distinct from those caused by mutation of the other known regulators of polarized BM protein secretion in the follicle cells, all of which cause BM proteins to accumulate on the apical surface (Denef et al., 2008; Devergne et al., 2014, 2017; Isabella and Horne-Badovinac, 2016; Lerner et al., 2013). Strong depletion using *Khc-73^3-3^ & Khc^RNAi^*, however, causes BM proteins to accumulate mainly on the apical surface. This result could indicate that, like Rab10, kinesins also play an earlier role in sorting BM proteins into a basolateral pathway. However, a second possibility is suggested by the observation that Rab10+ compartments also relocalize apically when kinesins are strongly depleted. BM proteins may still enter a Rab10+ compartment, but without plus end-directed kinesins present, a minus end-directed motor like dynein could redirect the vesicles apically. If true, this would imply that simply delivering BM protein vesicles to the apical membrane is sufficient for apical secretion in follicle cells, and that titrating the relative levels of plus and minus end-directed motors plays a key role in polarized secretion. New tools to visualize how BM proteins move through the secretory pathway will be needed to distinguish between these models.

It is likely that kinesin-based transport integrates with other cellular mechanisms in the follicle cells to ensure BM proteins assemble into a single sheet. Contributions from receptors for BM proteins like integrins and dystroglycan may play a role in determining where BM proteins assemble, and could aid in post-secretion movements of BM proteins such as those required for BM fibril formation (Campos et al., 2020; Jayadev and Sherwood, 2017). We previously showed that Col IV-encoding mRNAs are enriched basally in follicle cells and proposed that local Col IV protein synthesis and export through basally localized Golgis promotes polarized BM protein secretion (Lerner et al., 2013). We re-examined Col IV-encoding mRNAs in this study using a more sensitive detection method and detected more individual Col IV mRNAs distributed throughout the cell. Thus, kinesin-based transport of Rab10+ secretory vesicles may ensure that the subset of BM proteins that are synthesized away from the basal surface still reach the correct secretion site. Kinesin-based transport of BM proteins may be particularly important in mammalian cells, where a single, apically localized Golgi apparatus creates an even bigger spatial problem for BM protein secretion.

Finally, this work demonstrates that tightly controlling the sub-cellular site of BM protein secretion is critical for epithelial architecture. We previously showed that targeting BM protein secretion to lateral surfaces alters the architecture of the follicular BM in a way that is beneficial for the tissue - it allows the formation of the BM fibrils necessary for normal egg chamber elongation (Isabella and Horne-Badovinac, 2016). It is now clear, however, that BM protein secretion cannot occur anywhere along the lateral membrane, as secretion to the upper region of this domain causes BM protein networks to form between the cells. Moreover, when these ectopic networks attach to the main BM, they act as anchors that locally impede the collective migration of the follicle cells, distorting the cells’ shapes to such an extent that even the nuclei are deformed. We imagine that the formation of ectopic BM protein networks between cells would be detrimental to any epithelium undergoing cellular rearrangements. These would include processes like cell division and cell extrusion that are required for tissue growth and homeostasis, as well as processes like cell intercalation and apical constriction that underlie changes in tissue shape (Kozyrina et al., 2020). Given that BM assembly and epithelial morphogenesis often coincide during development (Jayadev and Sherwood, 2017; Walma and Yamada, 2020), we propose that kinesin-based transport of BM proteins toward the basal surface may provide a general mechanism to ensure that these two key aspects of epithelial development do not interfere with one another.

## Author Contributions

L.L. Z. and S.H-B. conceived the study. A.L.Z. designed experiments, generated new reagents, performed experiments, analyzed the data, and prepared the figures. A.L.Z. and S.H-B. wrote the manuscript.

## Acknowledgements

We thank Lynn Cooley, Robin Hiesinger, Nina Sherwood, Pejmun Haghighi, Dan Bergstralh, Vladimir Gelfand, and Chris Doe for *Drosophila* stocks. Younan Li for help with Sobel operator analysis and MATLAB code. Members of the S.H-B., Munro, Fehon, and Glick labs for experimental advice, and members of the S.H-B. lab, Kriza Sy, and Serapion Pyrpassopoulos for comments on the manuscript. Funding was provided by the following: American Heart Association 16POST2726018, American Cancer Society 132123-PF-18-025-01-CSM, and Chicago Biomedical Consortium FP064171-01-PR postdoctoral fellowships to A.L.Z. American Cancer Society RSG-14-176 and NIH R01 GM136961 to S.H-B.

## METHODS

### RESOURCE AVAILABILTY

#### Lead Contact

Further information and requests for resources and reagents should be directed to and will be fulfilled by the Lead Contact, Sally Horne-Badovinac.

#### Materials Availability

New *Drosophila* lines and plasmids generated in this study are available by request to the Lead Contact above.

#### Data and Code Availability

Sequence data is included in Supplemental Table 3. This study did not generate new code.

### EXPERIMENTAL MODEL AND SUBJECT DETAILS

#### Drosophila care

*Drosophila melanogaster* were reared on cornmeal molasses agar food at 25°C using standard techniques. The genotypes used in each experiment are listed in Supplemental Table 1. Females were aged on yeast with males prior to dissection; temperatures and yeasting conditions used for each experiment are in Supplemental Table 2 indexed by figure.

## METHOD DETAILS

### Generation of *Khc-73^3-3^* allele

Clustered regularly interspaced short palindromic repeats (CRISPR) genome editing was used to induce a lesion near the amino-terminus of Khc-73 within the motor domain. Two guide RNAs were selected using *Drosophila* RNAi Screening Center’s “Find CRISPRs” on-line tool, one within exon 3 (5’ <u>ATATGCACGCATTATAGCCC</u>**TGG** 3’) and one within exon 4 (5’ <u>CTTGTACATAAGCTCGGGTG</u>**TGG** 3’). The PAM motifs are in bold, and the underlined sequences were cloned into pU6-BbsI-chiRNA following the methods in (Gratz et al., 2013, 2014) and the website flycrispr.org. *GenetiVision* injected chiRNA plasmids into embryos expressing nanos-Cas9 from the x chromosome. Individual lines were established and screened by PCR and sequencing. The *Khc-73^3-3^* allele has a lesion near the guide site in exon3 only that results in a stop codon. The sequence of the resulting lesion, primers, and all plasmid sequences are available in Supplemental Table 3. We focus on developmental stages 7 and 8 in this mutant because there are not obvious defects in BM protein secretion earlier in development with loss of only Khc-73. BM protein secretion is highest at stage 7 which could make it easier to detect secretion defects, but we cannot rule out developmental changes in secretion regulation.

### Egg chamber dissections

Ovaries were removed from yeasted females using 1 set of Dumont #55 forceps and 1 set of Dumont #5 forceps in live cell imaging media in a spot plate (Schneider’s *Drosophila* medium containing 1X Penicillin-Streptomycin, 15% fetal bovine serum, and 200 μg/ml insulin). Ovariole strands were mechanically removed from muscle with forceps. Egg chambers older than stage 9 were cut away from the ovariole strand in the stalk region using a 27-gauge needle. For additional methods and videos of dissection, see (Cetera et al., 2016).

### Live imaging sample preparation

For live imaging, dissected ovarioles were quickly moved to a fresh well of live imaging media in a spot plate. In some live experiments, noted in the appropriate sections, CellMask^TM^ Orange or Deep Red plasma membrane stain was used to visualize cell edges and aid in imaging setup. Either version of CellMask^TM^ was added at 1:2000 to ovarioles in live imaging media for 15 min. The ovarioles were then washed 2× in fresh live imaging media to remove excess stain. To make a live imaging slide, 1-5 ovarioles were transferred to a glass slide in 10 µl of live imaging media. Glass beads (between 10 and 50) with a mean diameter of 51 µm were added for use as spacers, and arranged around the egg chambers using an eyelash tool. A 10 µl drop of fresh live imaging media was added to a #1.5 22×22 mm square cover glass to prevent bubbles, and slowly lowered onto the egg chambers. The slide was sealed with melted petroleum jelly before imaging. New dissections were done every hour to avoid artifacts arising from extended *ex vivo* culture.

### Extracellular stain of Col IV-GFP

Egg chambers were fixed for 6 min at room temperature (RT) in 4% EM grade formaldehyde in phosphate buffer saline (PBS), washed 3×5 min in PBS at RT, and stained with 1:2000 anti-GFP nanobody conjugated to Alexa Fluor^®^ 647 (GFP-Booster) and 4′,6-diamidino-2-phenylindole (DAPI) for 15 min at RT with rocking *without* permeabilization to allow the nanobody access to only the extracellular pool of Col IV-GFP. Samples were washed 3×5 min with PBS.

### Immunostaining

Egg chambers were fixed in 4% EM-grade formaldehyde in PBS with 0.1% Triton X-100 (PBST) for permeabilization for 15 min at room temperature, and then washed 3×10 min in PBST. For microtubule (MT) staining, 8% formaldehyde in PBST was used to better preserve MTs (Doerflinger et al., 2003). Egg chambers were incubated with primary antibodies diluted in PBST overnight at 4°C with rocking. Primary antibody dilutions: aPKC (1:100), Dlg (1:10), Tango1 (1:1000), GM130 (1:500), anti-acetylated ⍺-tubulin (1:100). Egg chambers were washed from primary antibody 3×10 min in PBST with rocking at RT. Secondary antibodies were diluted 1:500 in PBST and incubated with egg chambers for 3 hrs at RT with rocking, followed by washing 3×10 min in PBST.

### smiFISH

Single molecule inexpensive fluorescent in situ hybridization (smiFISH) was based on (Tsanov et al., 2016) and protocols provided by Matt Ronshaugen’s lab. In this technique, DNA probes specific to the mRNA of interest are fused to a “flap” sequence that will anneal to a complementary, fluorescently-blabeled “flap” sequence to allow visualization of the mRNA.

### Probe Design and Annealing

DNA probes specific to *col41a* mRNA (based on cDNA clone RE33133) were designed using LGC Biosearch Technologies’ Stellaris^®^ RNA FISH Probe Designer with the following settings: probe length 20 bases, masking level 5, minimal spacing 2 bases. 48 probes were ordered for *col4a1*. Each probe contained 20 nucleotides complementary to mRNA for *col4a1* followed by Flap-X (5’CCTCCTAAGTTTCGAGCTGGACTCAGTG 3’) appended to the 3’ end. Probes were ordered from Integrated DNA Technologies (IDT) as 100 µM stocks in Tris-EDTA, pH 8.0 (TE) in a 96-well plate.

Probes were mixed at equal molar ratios to make a stock of unlabeled probes and stored at −20°C. Working stocks were diluted 5-fold in TE buffer before use. Fluorescently labeled Flap-X binding probes were ordered from IDT as DNA oligos with 5’ and 3’ Cy5 modifications and resuspended in TE at 100 µM. All probe sequences are listed in Supplemental Table 3. *Col4a1* probes and fluorescent probes were annealed immediately before use by mixing: 2 µl of probe set, 0.5 µl of 100 µM Cy5-FlapX, 1 µl of New England Biolabs^®^ Buffer 3, and 6.5 µl water. A PCR machine was used to incubate mixtures at 85°C for 3 min, 65°C for 3 min, and 25°C for 5 min.

### Hybridization

Egg chambers were fixed in 4% EM-grade formaldehyde in PBST for 15 min at room temperature (RT) and washed 3×5 min in PBST. Egg chambers were exchanged into a 1:1 mixture of PBST and smiFISH wash buffer [5 ml 20X SSC (0.3M sodium citrate, 3M NaCl, pH7.0), 5 ml deionized formamide, 40 ml nuclease-free water] and incubated at RT for 10 min. Egg chambers were washed 2× in smiFISH wash buffer, followed by a final incubation of 30 min at 37°C. smiFISH hybridization buffer (1 g dextran sulfate, 1 ml 20x SSC, 1 ml deinionzed formamide, 7.5 ml nuclease-free water) was warmed to 37°C. Egg chambers were incubated with a mixture of 10 µl of annealed probes in 500 µl of smiFISH hybridization buffer at 37°C for 16 hrs protected from light. To wash, 500 µl of smiFISH wash buffer was added to dilute hybridization buffer and the egg chambers were spun briefly in a table-top micro-centrifuge to pellet them. Egg chambers were washed 3×10 min at 37°C in 500 µl of smiFISH wash buffer. This was followed by incubating at RT in a 1:1 dilution of smiFISH wash buffer with PBST, followed by one 10 min incubation in PBST containing DAPI and Alexa Fluor™ 488 Phalloidin, followed by 2×10 min washes with PBST at RT.

### Mounting fixed samples

For all types of fixed samples, the majority of buffer was removed from samples and they were mounted in ∼35 μl SlowFade^TM^ antifade or VECTASHIELD^Ⓡ^ on a slide with a #1.5 22×50 mm coverslip, sealed with nail polish, and stored at 4°C prior to imaging.

### Microscopy

#### Fixed and live laser scanning confocal imaging

Imaging was performed on a Zeiss LSM 800 laser scanning confocal microscope with Zen blue, 63× Plan A apochromat 1.4NA oil objective. Live imaging was performed at RT. Used in Figures 1A, 1D, 2A, 2C, 3A, 4A, 4C, 5A, 6A, 6C, 6F, 6H, 6I, 7A, 7C, 7E, S1B, S1D, S1F, S2A, S2B, S3A, S3F, S4A, S4B, S5A, S6A, S6B, Movie 1, and Movie 6.

#### Airyscan fixed imaging

Zeiss LSM 880 laser scanning confocal microscope with Airyscan and a 63× Plan A apochromat 1.4NA oil objective run by Zen black. Used in Figures 3D, 3E, and Movie 2.

#### Live spinning disk confocal and partial TIRF imaging

Imaging was performed on a Nikon Ti-E inverted microscope equipped with solid-state 50 mW, 481 and 561 nm Sapphire lasers (Coherent), a Yokogawa CSU-X1 spinning-disk scan head, and an Andor iXon3 897 electron-multiplying charged-coupled device (EM-CCD) cameras run by MetaMorph software. When using the TIRF microscope, we adjusted the laser below the critical angle to illuminate a thicker region of the sample (partial TIRF). Live imaging was performed at RT. Spinning disk used in Figure 3G and Movie 3. Partial TIRF used in Figures 3C, 4F, S3C, S5C, S6E, Movie 4, and Movie 5.

### Quantification of lateral BM proteins

Single confocal images were taken in a plane through the lateral domains, near the apical surfaces of follicle cells from late stage 8 egg chambers when new Col IV-GFP synthesis is low and ectopic lateral Col IV-GFP in mutant egg chambers is easily visible. The average intensity of Col IV-GFP associated with lateral domains was measured by segmenting the lateral domains based on anti-Dlg staining. In Fiji, Dlg images were processed as follows: background was subtracted with Rolling Ball Background Subtraction with a radius of 10 pixels, Threshold was used to create a binary mask, Despeckle was used to remove noise, Dilate was used to fill in gaps in the lateral edges, Skeletonize was used to reduce the mask to a one pixel outline, and finally Dilate was used to reach a uniform line thickness of 0.7 µm. This binary mask was used to measure the mean intensity of Col IV-GFP within only the lateral regions. In some images, the Dlg staining quality was too noisy for automatic segmentation, and the cell edges were manually traced in Fiji and converted to the same thickness as the automatic segmentation. For LanA-GFP and Perlecan-GFP, samples were counterstained with Alexa Fluor™ 647 Phalloidin, not anti-Dlg. F-actin highlights the cell edges, which were manually traced in Fiji and then converted to the same thickness as the automatic segmentation. The data were normalized to the mean of the control egg chambers for easy comparison as fold-change relative to control.

### Quantification of extracellular lateral Col IV-GFP

Additional processing was required for lateral extracellular Col IV-GFP because the “near basal” plane is near the BM, which creates high background fluorescence. Extracellular Col IV staining is necessary during stages when new Col IV protein synthesis is high (stage 7 in these experiments) and Col IV-GFP foci are scattered throughout the cytoplasm, obscuring the secreted, lateral Col IV-GFP population.

Three planes were chosen from a confocal z-stack of mosaic *Khc-73^3-3^* tissue for analysis: a plane “near basal”, or about 1 μm above the BM, a plane through the middle of the cell (“mid-cell”), and a plane just below the apical surface of cells (“near-apical”). The lateral cell edges were manually traced in Fiji with a line thickness of 1.12 μm at each z-plane. The mean intensity of the anti-GFP nanobody staining (extracellular Col IV-GFP, see Figure S1F) was measured in the segmented lateral regions for both the control and *Khc-73^3-3^* cell regions. The mean intensity was also measured in the inverse regions, within the cell centers where there should not be any extracellular staining; the mean intensity in these cell center regions was used for background subtraction in each plane, which is particularly important for the images near the basal surface where the out of focus fluorescence from the BM generates high background. The background corrected mean lateral intensity is calculated: mean intensity lateral regions – mean intensity central regions. To allow the data to be read as fold change between control and *Khc-73^3-3^* cells in one egg chamber, and be compared across z-planes, all data was divided by the mean of the control cells in the “near basal” images for each egg chamber.

### Line-scans of β-catenin and Col IV-GFP

Confocal images of cross-sections through *Khc-73^3-3^* egg chambers expressing Col IV-GFP and stained for anti-β-catenin were taken. In Fiji, lines of 0.58 µm thickness were manually drawn along 20 cell-cell junctions containing ectopic lateral Col IV-GFP, starting above the apical surface and extending along the lateral domain. The average intensities of Col IV-GFP and β-catenin were measured along the line-scan using Plot Profile and exported to Microsoft Excel. The maximum intensity of β-catenin was used to determine the location of the zonula adherens and used as a comparison point to align all the traces, where the zonula adherens was set to a distance of zero. All individual traces for Col IV-GFP intensity were plotted to show the range of accumulation patterns. Β-catenin is drawn as a line at zero as it is used only as a reference point for the zonula adherens junction that demarcates the lateral and apical domains.

### Colocalization of Col IV-GFP and the ER

Egg chambers expressing UAS-RFP-KDEL as a luminal marker of the ER and Col IV-GFP were dissected and stained with CellMask^TM^ Deep Red as described. Egg chambers were imaged live to better preserve ER structure. For the images shown, three consecutive frames from a time-lapse taken 1 sec apart were averaged to decrease noise.

### Quantification of YFP-Rab10

#### At basal trailing cell edges

Images were taken along the basal surface of cells in mosaic tissues. YFP-Rab10 is present as a diffuse cytoplasmic signal as well as a more intense signal on punctae and tubules at basal trailing cell edges, likely representing the vesicular structures of interest. To focus our measurements on the YFP-Rab10 associated with these putative vesicular structures, we quantified YFP-Rab10 intensity changes in only basal trailing cell edges so that the mean YFP-Rab10 measurements would not be dominated by the cytoplasmic signal that covers a much larger area of cells. In Fiji, a line with a thickness of 1.2 μm was manually drawn along the back of each cell and the mean YFP-Rab10 intensity was measured in each cell. To normalize the YFP-Rab10 intensities across egg chambers, the entire control cell area was manually segmented in Fiji and the mean YFP-Rab10 intensity measured. All individual cell values were divided by this value. In addition to individual cell measurements, we also calculated the mean of all cells per genotype for each egg chamber. Statistical tests were performed using these mean egg chamber values, but all individual cells are also plotted to show the variability in the underlying data (Lord et al., 2020).

#### Near the ERES

YFP-Rab10 is localized near ERES/Golgi regions throughout the volume of the cells. Near the basal surface, the tubules at the backs of cells are brighter than the signal by the Golgi, and their close, and sometimes overlapping, signal makes it difficult to measure the levels of YFP-Rab10 near ERES/Golgi.

In the middle of the cells, the majority of the YFP-Rab10 is concentrated near the ERES/Golgi so we chose this plane to allow easier segmentation. Tango1 staining at the ERES was used to segment these regions with the following steps: Rolling Ball Background Subtraction with radius 10 pixels was used, Threshold was used to create a binary mask of the ERES, and noise was removed by running Despeckle 2× in Fiji. The ERES was expanded using Dilate 2× so it would encompass the region adjacent to the ERES where Rab10 is normally found. The mean intensity of YFP-Rab10 within these ERES masks was measured in both the control cells and *Khc-73^3-3^* cells in each mosaic egg chamber, resulting in a single mean value per genotype per egg chamber. Since the same mosaic egg chambers were also used to measure the basal trailing edge YFP-Rab10 intensity, just at different z-planes, these ERES means were normalized using the same value used to normalize YFP-Rab10 at the basal trailing edge between egg chambers.

### Quantification of the effect of HA-Khc-73 OE

We used the *traffic jam-Gal4* (*tj-Gal4*) driver to express UAS-HA-Khc73 (Siegrist and Doe, 2005). The *tj-Gal4* driver usually expresses UAS transgenes in all follicle cells, but some UAS transgenes in our experience express in only a subset of cells, which we refer to as “patchy” expression in the text. We do not know why UAS-HA-Khc-73 in particular expresses in this patchy way. This phenomenon has also been described as variegation in expression in follicle cells, thought to stem from epigenetic changes (Lee and Spradling, 2014; Lee et al., 2017; Skora and Spradling, 2010). Since we can stain for the HA epitope tag, we were able to select non-expressing cells and HA-Khc-73 expressing cells. Images were taken near the basal surface of tissues patchily overexpressing HA-Khc-73, which clusters at the basal trailing edges of cells. To determine if HA-Khc-73 recruited other proteins to these clusters, we measured intensities in 1.2 μm lines drawn along the basal trailing edges of cells, similar to the procedure used to measure basal trailing edge YFP-Rab10. We chose to measure along the trailing edges instead of segmenting the HA+ regions because this more general location could be selected in the non-expressing cells as a control for baseline localization of candidate proteins to the trailing edge of cells. Cells that were negative for HA-Khc-73 staining (“control”) or positive for HA-Khc-73 were selected from within the same egg chamber. The mean intensity of the relevant endogenously tagged, fluorescently labeled protein (YFP-Rab10, Col IV-GFP, PDI-GFP (ER), or anti-GM130 (cis Golgi) staining) was measured for each cell. For each egg chamber, all individual cell measurements were normalized by dividing by the mean value for control cells within that egg chamber. In addition to individual cell measurements, we also calculated the mean of all cells per group (“control” or “HA-Khc-73”) for each egg chamber. Statistical tests were performed using these mean group values, but all individual cells are also plotted to show the variability in the underlying data (Lord et al., 2020).

### Live imaging YFP-Rab10 in HA-Khc-73 OE cells

We used the *traffic jam-Gal4* (*tj-Gal4*) driver to express UAS-HA-Khc73 and UAS-YFP-Rab10. Images were collected in a continuous 200 ms stream to follow the rapid movements of YFP-Rab10, using partial TIRF.

### Live imaging and tracking of YFP-Rab10

#### Selection of egg chambers

In some experiments, CellMask^TM^ Orange plasma membrane stain was used to visualize cell edges. To ensure egg chambers were not damaged during dissection or sample preparation, we first determined if they were migrating at a normal rate. A 5-30 min time-lapse using 300 ms exposures taken every 15 sec was taken for each egg chamber to ensure it was migrating at a normal speed (a cutoff of > 0.4 μm/min was set for the stage 7 egg chambers used). Sets of control and *Khc-73^3-3^* egg chambers were always imaged in the same session to prevent biases from day-to-day variability in live imaging conditions. Since partial TIRF imaging was used, there is uneven illumination across the field of cells.

#### Tracking and analysis

To allow tracking of rapidly moving individual YFP-Rab10 punctae, time-lapses with 300 ms exposures captured every 1 sec for 2 min were made. YFP-Rab10 punctae were manually tracked using the Manual Tracking plugin in Fiji. All punctae that moved at least 3 pixels (0.3 μm) per 1 sec time-step over at least 3 consecutive frames were tracked in at least 5 cells per egg chamber to obtain a minimum of 50 puncta tracks per egg chamber, in 5 egg chambers per genotype. We sometimes observed a change in direction of a YFP-Rab10 puncta. Each segment in a single direction was counted as a separate “run”, allowing a single puncta to represent multiple runs. Each run was scored by eye for its overall direction, either towards the trailing or leading edge of the cell. For each egg chamber, the number of tracks moving towards the trailing edge of the cell was divided by the number of tracks moving towards the leading edge of each cell, such that a value greater than 1 indicates a bias in movement towards the trailing edge where YFP-Rab10 accumulates. In addition, the speed of each “run” was calculated. Speed was calculated as the Euclidian frame-to-frame displacement. All individual “run” speeds across all 5 egg chambers were plotted to compare the distributions between control and *Khc-73^3-3^* cells, and the mean speed per egg chamber was also plotted and used for statistical comparison between genotypes. All analysis was done on original images; for display in figures, images were rotated in Fiji so that the direction of cell migration is toward the bottom of the page.

#### Editing of images for *Movie 5*

For movies, images were rotated such that the direction of cell migration is toward the bottom of the page. The CellMask staining channel was processed with a Rolling Ball Background subtraction, radius 20 in Fiji. Both the YFP-Rab10 and CellMask channels were processed with Bleach Correction in Fiji for the first segment of Movie 5. The images of trajectories were generated in Fiji with the Manual Tracking plug-in.

### Quantification of apical and basal YFP-Rab10 levels

In images of cross-sections through egg chambers, lines of 0.6 μm thickness were manually drawn along the basal surface and apical surface of the follicular epithelium of each egg chamber and the mean intensity of YFP-Rab10 measured in Fiji. The ratio of the apical and basal surface values was calculated for each egg chamber, such that a ratio greater than 1 indicates an enrichment of YFP-Rab10 on the apical surface, and a ratio less than 1 indicates an enrichment of YFP-Rab10 on the basal surface.

### Measurement of fraction of Col IV within fibrils in the BM

As a way to quantify changes in the organization of Col IV-GFP within the BM, we measured the fraction of Col IV-GFP intensity associated with fibril-like structures in the BMs of different genotypes. We have previously performed this analysis and refer to it as “fibril fraction” (Isabella and Horne-Badovinac, 2016). First, we determined if the overall mean intensity of Col IV-GFP was the same.

The mean intensity of Col IV-GFP in a confocal plane though the BM was measured in a square region (3600 μm^2^) for each egg chamber in Fiji. The genotype that causes apical secretion had significantly reduced Col IV-GFP in the BM, so it was excluded from fibril fraction analysis. To segment the fibrils, the median intensity of Col IV-GFP was measured for each egg chamber and a threshold of 1.35x median intensity was used to define the brighter, “fibril” areas. The Col IV-GFP intensity in these “fibril” regions was divided by the total Col IV-GFP intensity in the image to obtain the % of total Col IV-GFP intensity associated with fibrils.

### Color 3D projections

#### Of MTs

The temporal color-code plug-in in Fiji was used to generate a color z-projection of a 1.8 µm Airyscan confocal z-stack of MTs in a wild type egg chamber. An antibody specific to acetylated MTs was used for this imaging because this antibody has been used in all previous work characterizing the basal MT array in follicle cells (Aurich and Dahmann, 2016; Chen et al., 2016; Viktorinová and Dahmann, 2013) and we wanted to ensure we visualized the same MT array as past work. Live imaging of MTs labeled with UAS-ChRFP-Tubulin or Jupiter-GFP show a similar organization of MTs at the basal surface (Figures 3G, S3C, S5C, and S6E).

#### Of intercellular BM protein network

The temporal color-code plug-in in Fiji was used to generate a color z-projection of Col IV-GFP in a confocal z-stack through the full-thickness of the follicle cells (9 µm). The cell outlines are Dlg staining of lateral cell edges in a single plane through the middle of FCs. The nuclei are maximum intensity projections of the DAPI signal from the z-stack to show the full shape of the nucleus. In the more apical slices of the z-stack, the large nuclei from the germline nurse cells were visible; these were manually circled and deleted from the Z-stack prior to making the maximum intensity projection. The nuclei highlighted in green with deformed/elongated shapes were manually highlighted in Illustrator.

### 3D projections of intercellular Col IV-GFP

A 3D confocal z-stack through the full thickness of the follicle cells was collected. The slices containing the BM were deleted to provide better contrast to visualize the intercellular Col IV-GFP network. 3D projections were generated in Fiji using the Plugin ClearVolume.

### smiFISH quantification

The density of *col4a1* mRNA precluded single-molecule counting. We used the intensity of the smiFISH probes as a proxy for mRNA levels. Three z-planes through the lateral domains of the follicle cells were chosen for analysis in each egg chamber to determine if the localization of mRNA changed in mutant cells: a plane along the basal surface, a plane through the mid-section of the cells, and a plane near the apical surface. Within the mosaic tissue, regions encompassing cells of different genotypes were manually drawn based on the nuclear clone marker in Fiji. The mean intensity of *col4a1* mRNA was measured for each genotype at each plane. Intensities between genotypes within mosaic tissues were compared within the same egg chamber at each plane by: dividing *Khc-73^3-3^* cells by control cells, or dividing *Khc^RNAi^* & *Khc-73^3-3^* cells by *Khc^RNAi^* only cells. The direction of follicle cell migration (to orient images) was determined by the organization of F-actin rich leading edges along the basal surface visualized by counter-staining egg chambers with Alexa Fluor™ 488 Phalloidin.

### Quantification of Patronin-GFP

Egg chambers were fixed and stained with Alexa Fluor™ 647 Phalloidin and DAPI. Confocal cross-sections were taken through egg chambers. In Fiji, a 1.1 μm thick line was manually drawn along the apical surface where Patronin is normally enriched (Khanal et al., 2016; Nashchekin et al., 2016), using the F-actin stain to visualize cell outlines. The mean apical intensity of Ubi-PatroninA-GFP in *Khc-73^3-3^* cells was divided by that of control cells in each egg chamber, resulting in a ratio of 1 when there is no change.

### Quantification of Jupiter-GFP

#### Intensity in cross-section

Confocal cross-sections were taken through live egg chambers stained with CellMask^TM^ Deep Red plasma membrane stain to aid in setting up imaging and identifying the borders of cells. In Fiji, the control and *Khc-73^3-3^* cells were manually outlined based on the clone marker, and the mean intensity of endogenously-tagged Jupiter-GFP (a MT-associated protein) was measured as a proxy for MT mass. The mean Jupiter-GFP intensity in *Khc-73^3-3^* cells was divided by that of the control cells in each egg chamber.

#### Basal MT alignment

Partial TIRF images of the basal surface of live mosaic egg chambers expressing Jupiter-GFP were taken. First, a 5-30 min time-lapse was collected to determine the direction of migration and to exclude non-migratory egg chambers damaged during dissection. To quantify MT alignment, we used an approach based on the Sobel operator (Gonzalez and Woods, 2017) to identify the sharp local gradients in fluorescent intensity present orthogonal to linear structures like MTs, as implemented in (Li and Munro, 2020). Images of Jupiter-GFP were convolved with the following 3×3 kernel Sobel operators to measure the x and y components of the fluorescent intensity gradient (Gx and Gy, respectively).

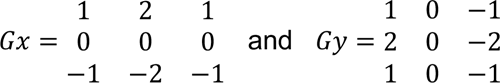

For each pixel, the angle and magnitude of the gradient can be calculated using the x and y components of the gradient. Since the gradient is orthogonal to the angle of the MT, the angle of the MTs θ is

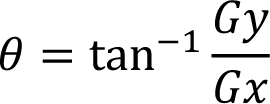

And the magnitude of the gradient, G, is 

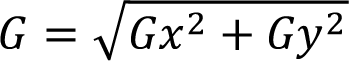

To create MT alignment distributions, regions of cells with different genotypes were manually outlined in Fiji based on the clone marker. The pixels within regions of a given genotype were binned by angle with weight G, collapsed to 0°-90° since we do not take into account angle relative to the A-P axis, and then normalized to 100%. The mean ± SD at each angle bin of all egg chambers of a given genotype were plotted on a rose diagram, where perfect alignment with the direction of migration would be 90° and alignment orthogonal to migration would be 0°. To statistically compare the amount of alignment between genotypes, the % of the alignment histogram within angles 60°-90° was used as proxy measurement of the population of “highly aligned MTs”. This “% highly aligned MTs” measurement was performed for each genotype in each egg chamber. All analysis was done on original images; for display in figures, images were rotated so that the direction of cell migration is toward the bottom of the page. MT intensity appears uneven across the egg chamber because these images were collected with partial TIRF which creates an uneven illumination depth and some interference patterns across the imaging field.

### Generation of Movies

Fiji was used to add labels and export movies as .avi, which were then converted to .mp4 using HandBrake.

### QUANTIFICATION AND STATISTICAL ANALYSIS

Egg chambers with visible damage from dissection or dying cells, or not migrating at a normal speed in live imaging experiments, were excluded. All experiments were replicated at least once. All statistical tests were performed in Prism8 or Prism9. MATLAB^®^ and Microsoft Excel were used as indicated in Methods for some data analysis. All data was tested for normality using the Shapiro-Wilks test and a non-parametric statistical test was chosen if a dataset was not normal. For comparisons between cells with different genetic perturbations within a mosaic tissue, a paired statistical test was chosen. When experiments compared data taken from different egg chambers, unpaired statistical tests were used.

ANOVA followed by a multiple comparisons test was used for comparison of more than two datasets. Many experiments compare the ratio between mutant and control cells, which would result in a value of 1 if there was no difference between groups; in these experiments a one-sample t test or the non-parametric Wilcoxon signed rank test was used to compare the experimental ratio to the theoretical value of 1. The number of biological replicates (n), specific statistical tests performed, and significance for each experiment can be found in the figure legends.

## RESOURCE TABLE

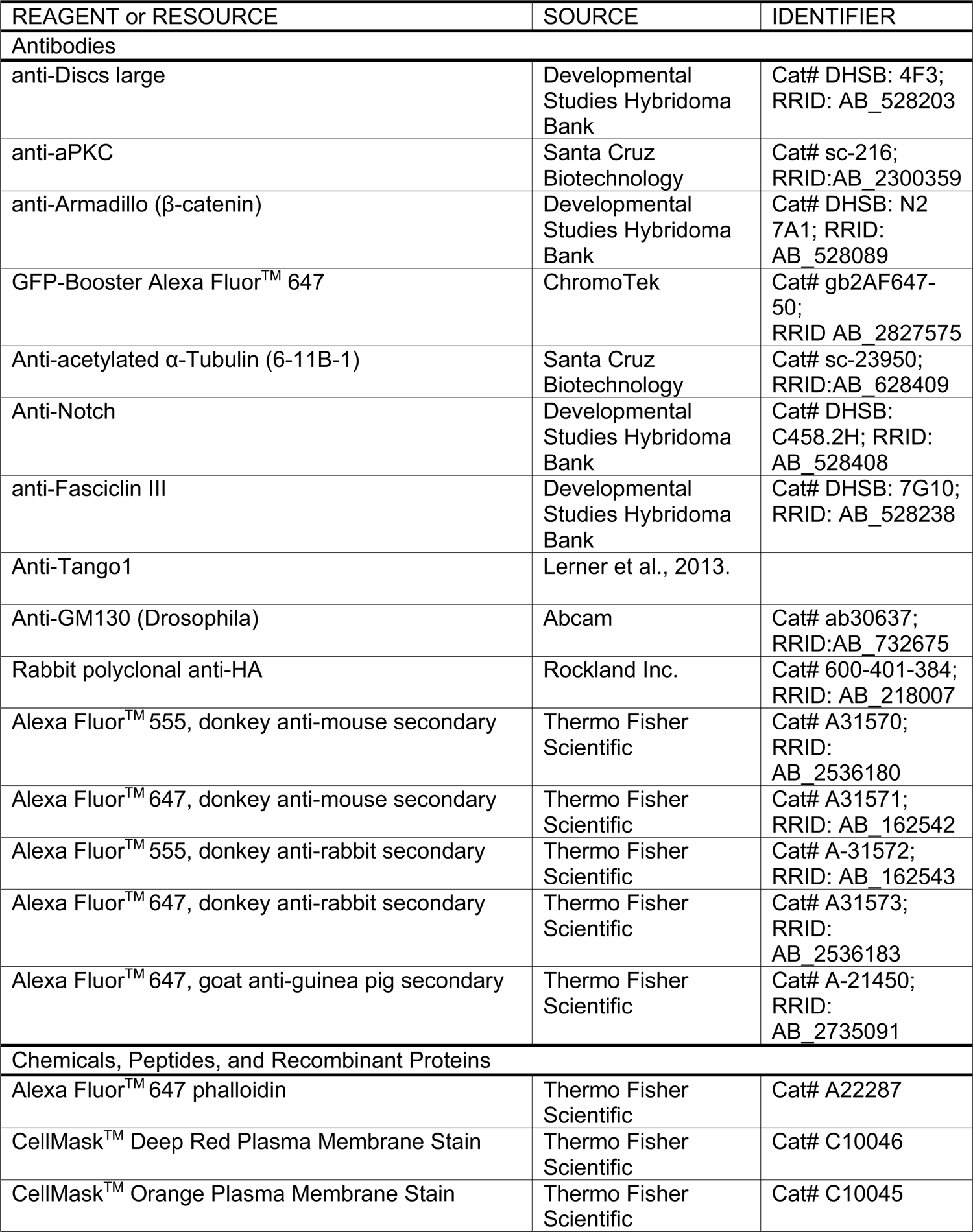
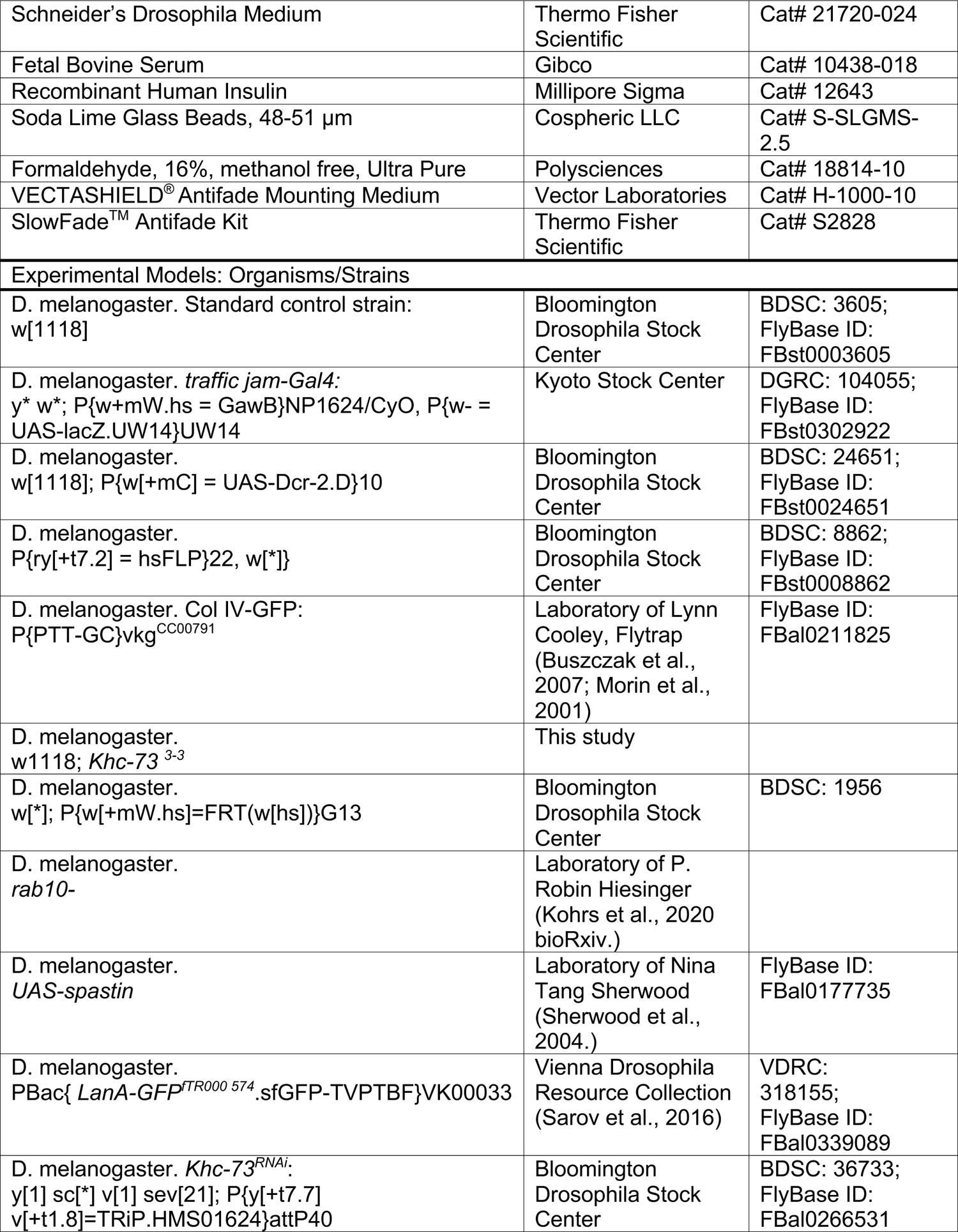
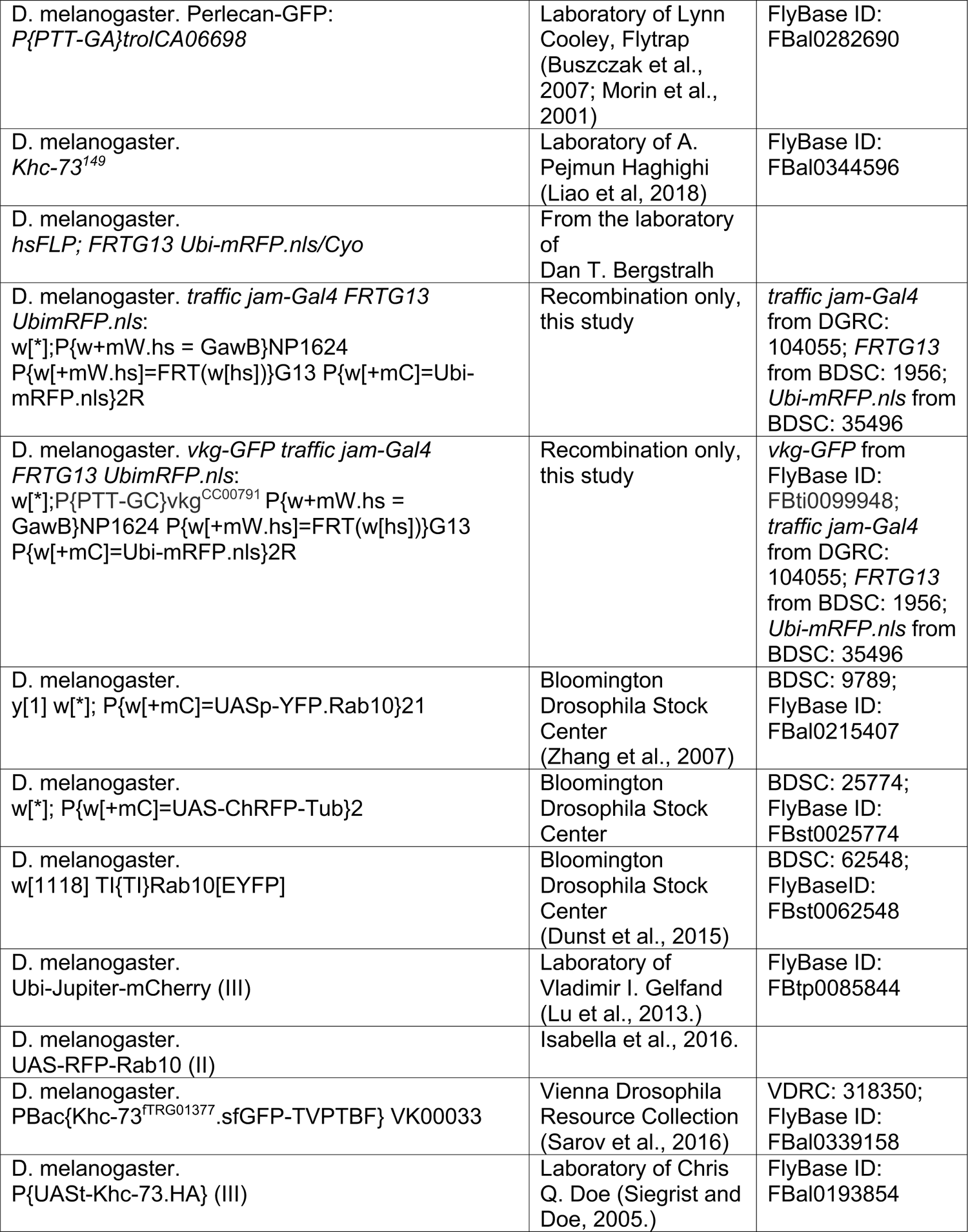
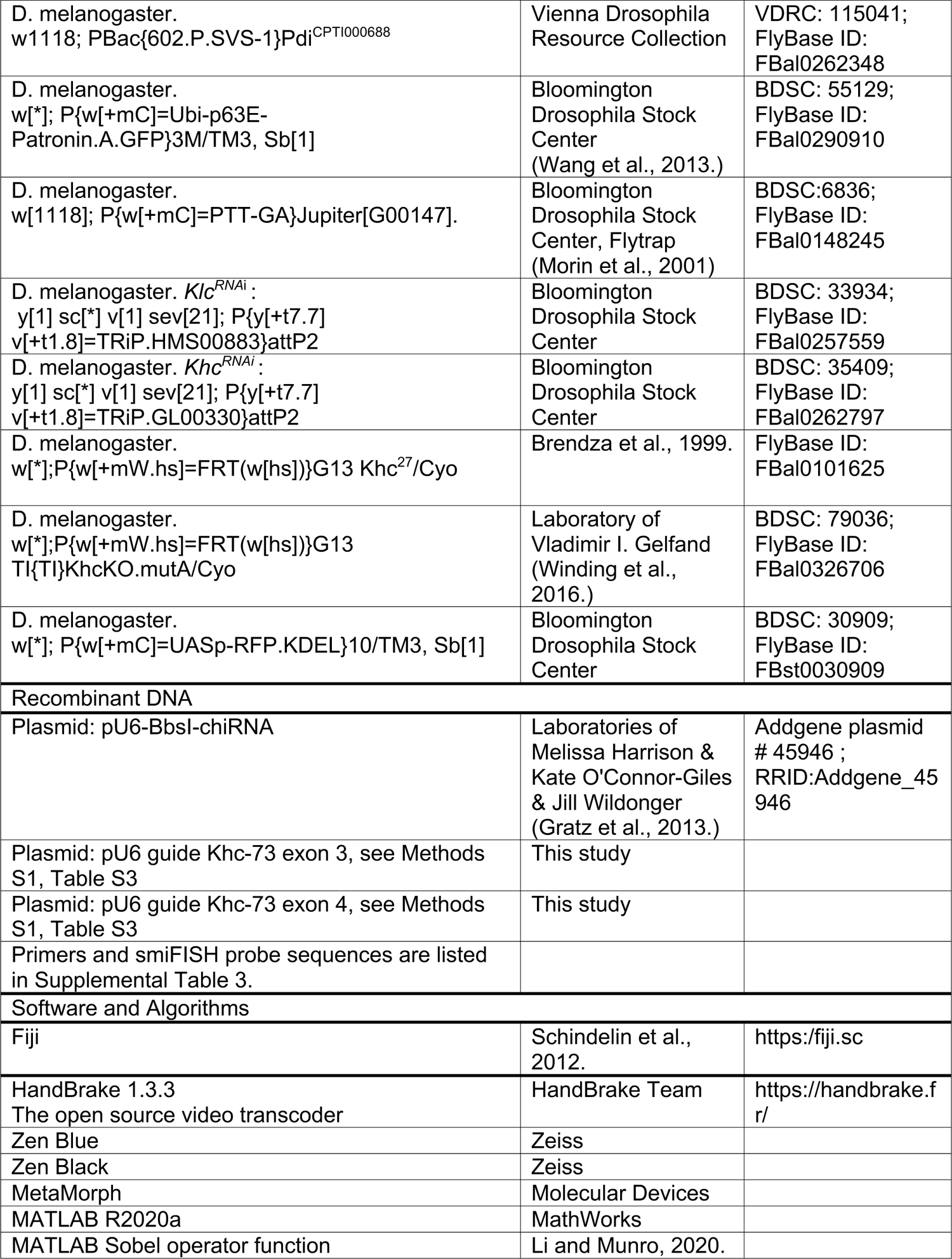
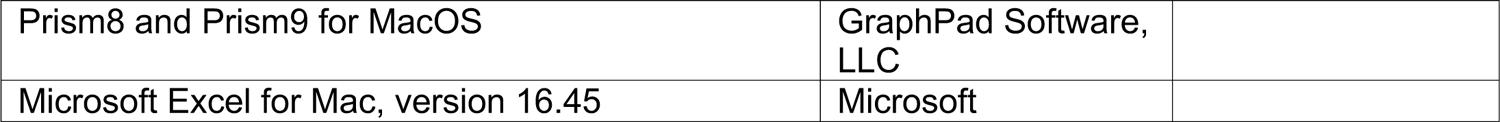

## SUPPLEMENTAL ITEMS LEGENDS

**Supplemental Table 3.** Sequences of reagents used for CRISPR of Khc-73, *Khc-73^3-3^* allele, and smiFISH probes, related to Figure S1B and Method Details. Supplemental excel sheet

**Movie 1. Background information on follicle cell migration and BM fibril deposition** Section 1 shows a confocal time-lapse taken along the basal surface of follicle cells in a stage 7 egg chamber. The BM is visualized with Col IV-GFP and cell edges are stained with CellMask. Follicle cells collectively migrate along the static BM (Haigo and Bilder, 2011). Section 2 diagrams how newly secreted BM proteins form fibrils in the space between cells, which then attach and are pulled on the BM as cells migrate (Isabella and Horne-Badovinac, 2016). Section 3 shows a time-lapse of fibril deposition. The BM is visualized with Col IV-GFP and cell edges are stained with CellMask. The BM has been bleached to facilitate visualization of newly deposited fibrils. Fibrils are manually highlighted with colored lines. As shown in Isabella and Horne-Badovinac, 2016).

**Movie 2. 3D organization of MTS** Images from an airyscan confocal z-stack of MTs (anti-acetylated ⍺-tubulin staining) in wild-type follicle cells. Inferno lookup table (LUT) represents intensity of MT staining. Movie starts at the basal surface and steps through the full-thickness of the epithelium. This is followed by repeating this stack with a focus on a single cell. The yellow box highlights the leading edge of the cell where MT bundles bend and integrate with the apical-basal MT array. A 3D projection of this yellow-boxed region follows the image stack. Stage 7 wild-type egg chamber. Scale bar, 10 µm.

**Movie 3. YFP-Rab10+ compartments move along MTs at the basal surface of follicle cells** Spinning disk confocal time-lapse of UAS-YFP-Rab10 and UAS-ChRFP-Tubulin dynamics along the basal surface of follicle cells. YFP-Rab10+ structures move rapidly along MTs and concentrate at the basal trailing edge of cells. Near the leading edge of cells, MTs appear to fade from view. Motile YFP-Rab10+ punctae appear and disappear in these regions. Stage 8 egg chamber. The tissue is also collectively migrating towards the bottom of the screen. Scale bar, 10 µm.

**Movie 4. Dynamic YFP-Rab10+ structures move into and out of HA-Khc-73-induced foci** Partial TIRF time-lapse of UAS-YFP-Rab10 in cells also overexpressing UAS-HA-Khc-73 taken along the basal surface of follicle cells. Relative intensity of the YFP-Rab10 signal is displayed using the inferno lookup table (LUT). A continuous 200 ms stream of images was collected to follow the rapid tubulovesicular dynamics in and out of the ectopic foci of YFP-Rab10 induced by overexpression of HA-Khc-73. The tissue is collectively migrating towards the bottom of the screen, but moves little over this time scale. Stage 7 egg chamber. Scale bar, 5 µm.

**Movie 5. Dynamics of YFP-Rab10+ compartments at the basal surface of control and *Khc-73^3-3^* follicle cells**

*Section 1*

Partial TIRF time-lapse of UAS-YFP-Rab10 with the cell edges labeled with CellMask. Images are taken every 15 sec to visualize how YFP-Rab10+ compartments are dynamically maintained at basal trailing edges as cells migrate. YFP-Rab10+ compartments are reduced at the basal surface of *Khc-73^3-3^* cells in comparison to controls. However, *Khc-73^3-3^* cells still have a smaller population YFP-Rab10+ compartments that appear at the basal surface and collect near the basal trailing cell edges. Stage 7 egg chambers. Scale bar, 10 µm.

*Section 2*

Partial TIRF time-lapse of UAS-YFP-Rab10 in the same egg chambers imaged every 1 second to allow tracking of individual YFP-Rab10+ puncta movements. YFP-Rab10+ punctae move rapidly towards both the trailing and leading edges of cells. Although there is less overall YFP-Rab10+ compartment accumulation at the basal surface of *Khc-73^3-3^* cells, the punctae present do move rapidly. Stage 7 egg chambers. Scale bar, 10 µm.

*Section 3*

Partial TIRF time-lapse of UAS-YFP-Rab10 along the basal surface. Images are taken every 1 sec. Two example cells were selected to show Rab10+ puncta trajectories. Trajectories are overlaid in the left panel and the original image is shown on the right. Cells are outlined in yellow at beginning of movie for orientation. The control time-lapse is followed by the *Khc-73^3-3^* time-lapse. Stage 7 egg chambers. Scale bars, 5 µm.

**Movie 6. BM cables act as anchors that impede cell migration, deforming cells and nuclei** A confocal z-stack beginning at the basal surface shows that the Col IV-GFP in a control egg chamber is mainly found in the plane at the basal surface. Follicle cells are hexagonally packed (anti-Dlg) and nuclei (DAPI) are round. In the *Khc-73^3-3^* egg chamber, a region with several cables of Col IV-GFP running from the basal BM to the apical surface deforms cell edges and nuclei. Stage 8 egg chamber. Scale bar, 5 µm.

