## Supplemental Data for "Kinesin-3 and kinesin-1 motors direct basement membrane protein secretion to a basal sub-region of the basolateral plasma membrane in epithelial cells"

### Figure S1

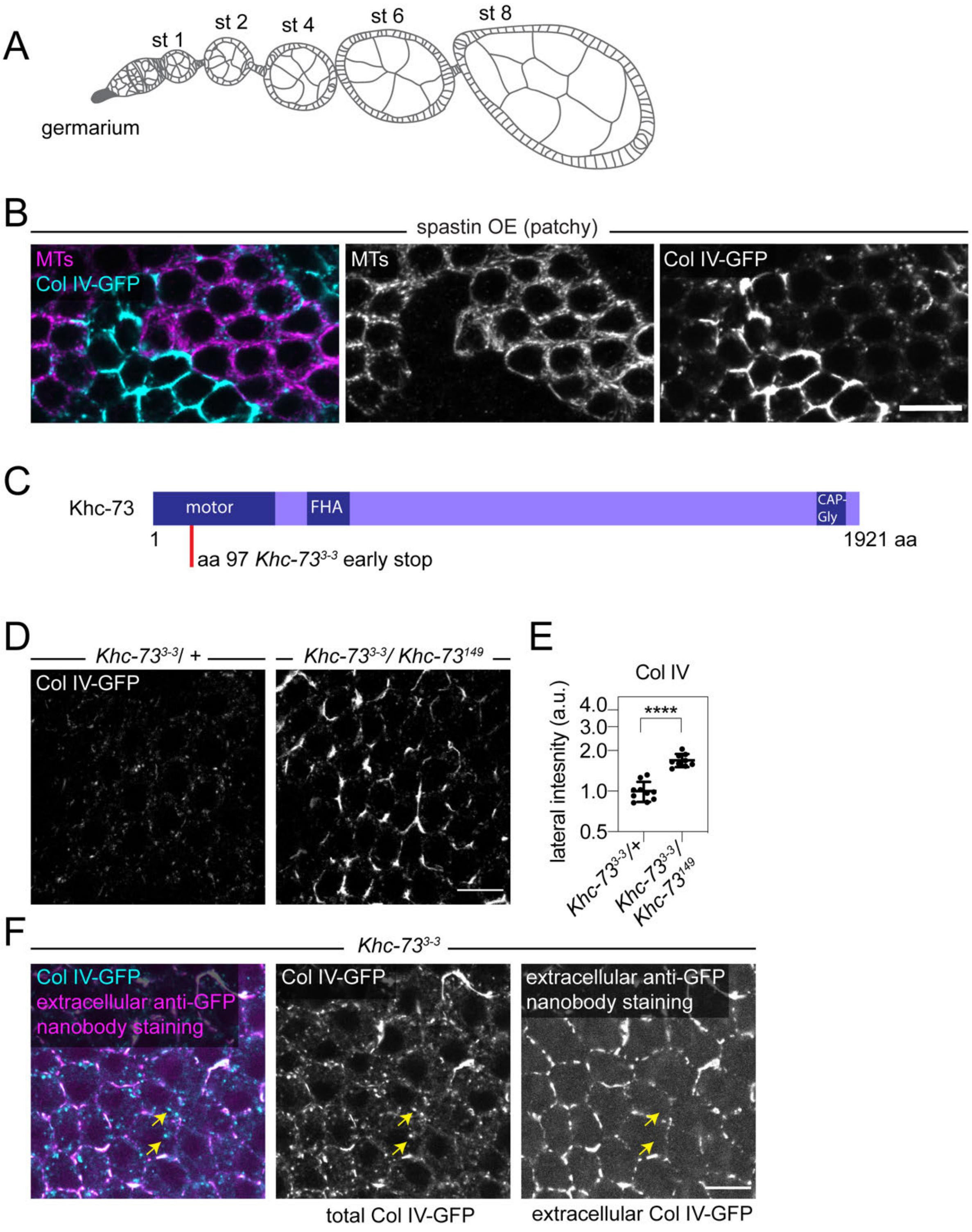
A
st 1
st 2
st 4
st 6
st 8
germarium
B
spastin OE (patchy)
MTs
Col IV-GFP
MTs
Col IV-GFP
C
Khc-73
motor
FHA
CAP-Gly
1
1921 aa
aa 97 Khc-73^3-3 early stop
D
Khc-73^3-3/ +
Khc-73^3-3/ Khc-73^149
Col IV-GFP
E
Col IV
lateral intesnity (a.u.)
4.0
3.0
2.0
1.0
0.5
****
Khc-73^3-3/+
Khc-73^3-3/ Khc-73^149
F
Khc-73^3-3
Col IV-GFP
extracellular anti-GFP nanobody staining
Col IV-GFP
extracellular anti-GFP nanobody staining
total Col IV-GFP
extracellular Col IV-GFP

**Figure S1, related to Figures 1 and 2, and Supplemental Table 3**

A. Illustration of an ovariole strand containing an array of egg chambers at different developmental stages. The germarium contains stem cell populations that generate new egg chambers which grow in size and elongate through 14 developmental stages to form an egg. Only stages 1-8 are shown here since we focus on stage 7 and 8 egg chambers in this study. Migration occurs between stages 2 and 8, and BM secretion occurs through stage 8.

B. Images showing Col IV-GFP accumulates at lateral cell edges in regions where MTs have been depleted by patchily overexpressing spastin. MTs visualized with anti-acetylated $\alpha$-tubulin staining.

C. Diagram of Khc-73 protein domains with the location of the lesion in *Khc-73*$^{3\text{-}3}$ indicated by red bar. See also Supplemental Table 3 for sequence.

D. Images showing lateral Col IV-GFP accumulation in a *Khc-73*$^{3\text{-}3}$/*Khc-73*$^{149}$ transheterozygous epithelium.

E. Quantification of increased lateral Col IV-GFP accumulation in *Khc-73*$^{3\text{-}3}$/*Khc-73*$^{149}$ epithelia. Data represent mean ± SD plotted on a log scale. Unpaired t test ****p<0.0001. In the order on graph, n= 10, 9 egg chambers.

F. Images of non-permeabilized *Khc-73*$^{3\text{-}3}$ tissue stained with an anti-GFP nanobody conjugated to Alexa Flour® 647 to highlight only extracellular Col IV-GFP. Yellow arrows indicate intracellular Col IV-GFP punctae not detected by the extracellular nanobody stain.

Stage 8 egg chambers (B and D). Stage 7 egg chamber (F). Scale bars, 10 μm.

### Figure S2

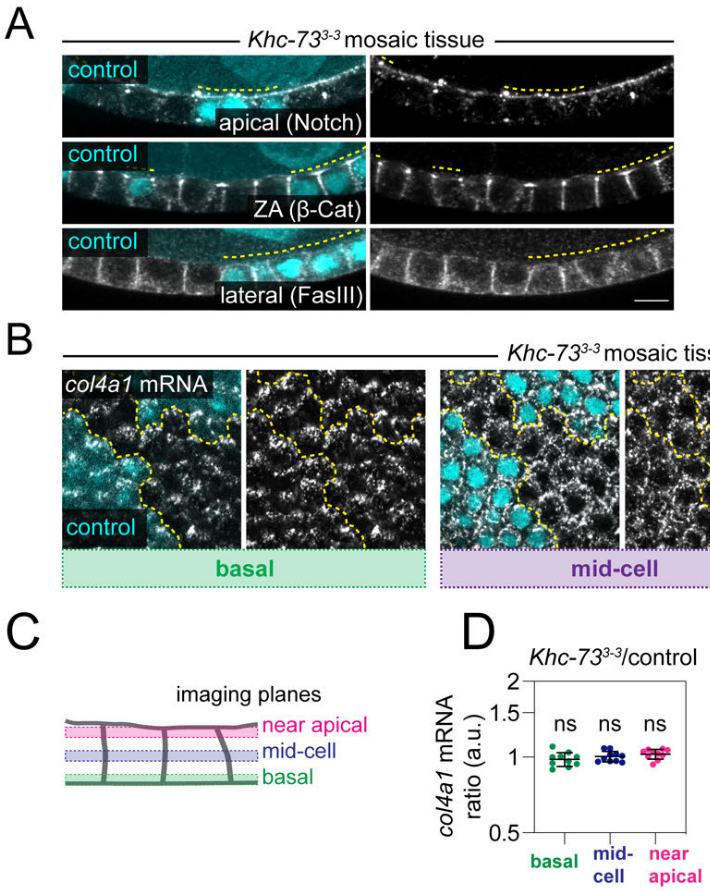
A
Khc-73^3-3 mosaic tissue
control
apical (Notch)
control
ZA (β-Cat)
control
lateral (FasIII)
B
Khc-73^3-3 mosaic tissue
col4a1 mRNA
control
basal
mid-cell
near apical
C
imaging planes
near apical
mid-cell
basal
D
Khc-73^3-3/control
col4a1 mRNA ratio (a.u.)
2
1.5
1
0.5
ns
ns
ns
basal
mid-cell
near apical

23

**Figure S2, related to Figure 1**

A. Images showing normal localization of transmembrane proteins to apical (Notch) or lateral (FasIII) domains, and β-catenin to the zonula adherens (ZA) in *Khc-73*$^{3\text{-}3}$ cells within a mosaic epithelium. All proteins detected by immunostaining. Control cells are marked in cyan and by a yellow dotted line drawn above them.

B. Images of single molecule inexpensive fluorescence in situ hybridization (smiFISH) for *col4a1* mRNA in *Khc-73*$^{3\text{-}3}$ mosaic tissue taken at 3 different z-planes through the lateral domains (see diagram in C). The dotted yellow line demarcates control and *Khc-73*$^{3\text{-}3}$ cells. Images are oriented so that migration is towards the bottom of the page.

C. Illustration of imaging planes in (B).

D. Quantification of *col4a1* mRNA intensity in *Khc-73*$^{3\text{-}3}$ cells relative to control cells in the same mosaic tissue at three different z-planes. The *col4a1* mRNA distribution along the apical-basal axis does not change in *Khc-73*$^{3\text{-}3}$ cells. Data represent mean ± SD plotted on a log scale. One sample t tests compared to the theoretical ratio of 1, ns $p > 0.05$. $n =$ 10 egg chambers.

Stage 7 egg chambers, except A (FasIII staining) is stage 6. Scale bars, 10 µm.

### Figure S3

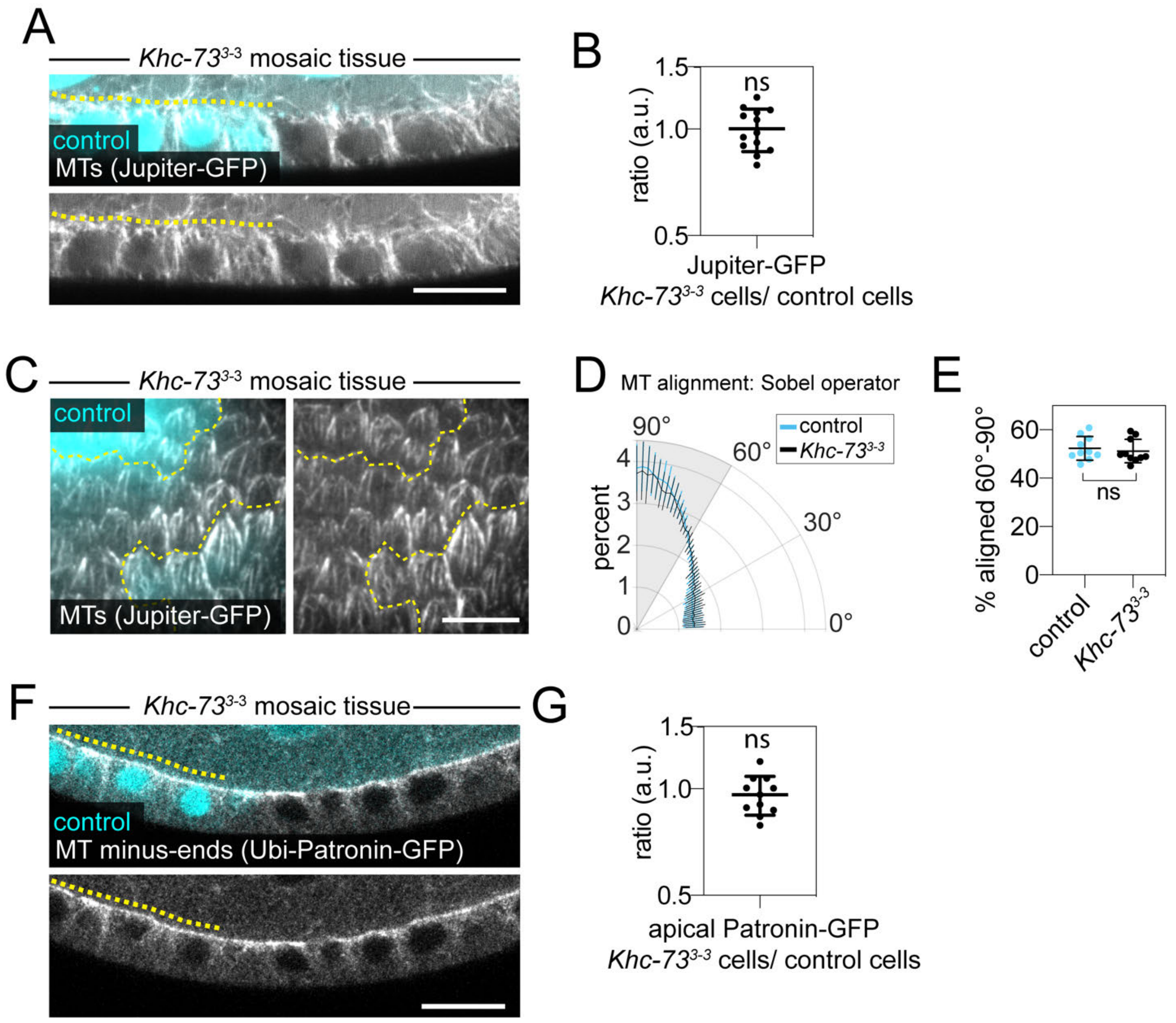
A
Khc-73$^{3-3}$ mosaic tissue
control
MTs (Jupiter-GFP)
B
ns
ratio (a.u.)
1.5
1.0
0.5
Jupiter-GFP
Khc-73$^{3-3}$ cells/ control cells
C
Khc-73$^{3-3}$ mosaic tissue
control
MTs (Jupiter-GFP)
D
MT alignment: Sobel operator
control
Khc-73$^{3-3}$
percent
90°
60°
30°
0°
E
% aligned 60°-90°
ns
control
Khc-73$^{3-3}$
F
Khc-73$^{3-3}$ mosaic tissue
control
MT minus-ends (Ubi-Patronin-GFP)
G
ns
ratio (a.u.)
apical Patronin-GFP
Khc-73$^{3-3}$ cells/ control cells

**Figure S3, related to Figure 1**

A. Image (live confocal) showing no change in MTs (Jupiter-GFP, an endogenously labeled MT-associated protein) running from apical to basal surfaces in *Khc-73*$^{3-3}$ cells within a mosaic tissue. Control cells are marked in cyan and by a yellow dotted line drawn above them.

B. Quantification of mean Jupiter-GFP intensity within follicle cells as a proxy for MT mass. The ratio of the Jupiter-GFP intensity in *Khc-73*$^{3-3}$ cells relative to control cells within the same mosaic tissue was calculated for each egg chamber. One sample t test compared to the theoretical ratio of 1, ns p>0.05. n=13 egg chambers.

C. Image from a live partial TIRF time-lapse of MTs (Jupiter-GFP) along the basal surface of a *Khc-73*$^{3-3}$ mosaic epithelium. MTs remain aligned along the basal surface of *Khc-73*$^{3-3}$ cells. The dotted yellow line demarcates control and *Khc-73*$^{3-3}$ cells. Images are oriented such that migration is towards the bottom of the page. MT intensity is uneven due to partial TIRF illumination.

D. Quantification of MT alignment using a Sobel operator shows the degree of MT alignment in control and *Khc-73*$^{3-3}$ cells from (C). The direction of cell migration is 90º. n=10 egg chambers.

E. Quantification of the enrichment in MT alignment in control and *Khc-73*$^{3-3}$ cells. To compare how much alignment with the direction of migration occurs in both genotypes, we measured the population aligned between 60º-90º from the Sobel operator (grey shading in D), where perfect alignment with the direction of migration is 90º. Paired t test, ns p>0.05. n=10 egg chambers.

F. Image showing the MT minus-end-binding protein Patronin (Ubi-Patronin-GFP) remains concentrated apically in *Khc-73*$^{3-3}$ cells. Control cells are marked in cyan and by a yellow dotted line drawn above them.

G. Quantification of Ubi-Patronin-GFP intensity in a line drawn along the apical surface, followed by calculating the ratio of the intensity in *Khc-73*$^{3-3}$ cells relative to control cells within the same mosaic epithelium. One sample t test compared to the theoretical ratio of 1, ns p>0.05. n=11 egg chambers.

Stage 7 egg chambers. Data represent mean ± SD. Data are plotted on a log scale in (B, G). Scale bars, 10 µm.

### Figure S4

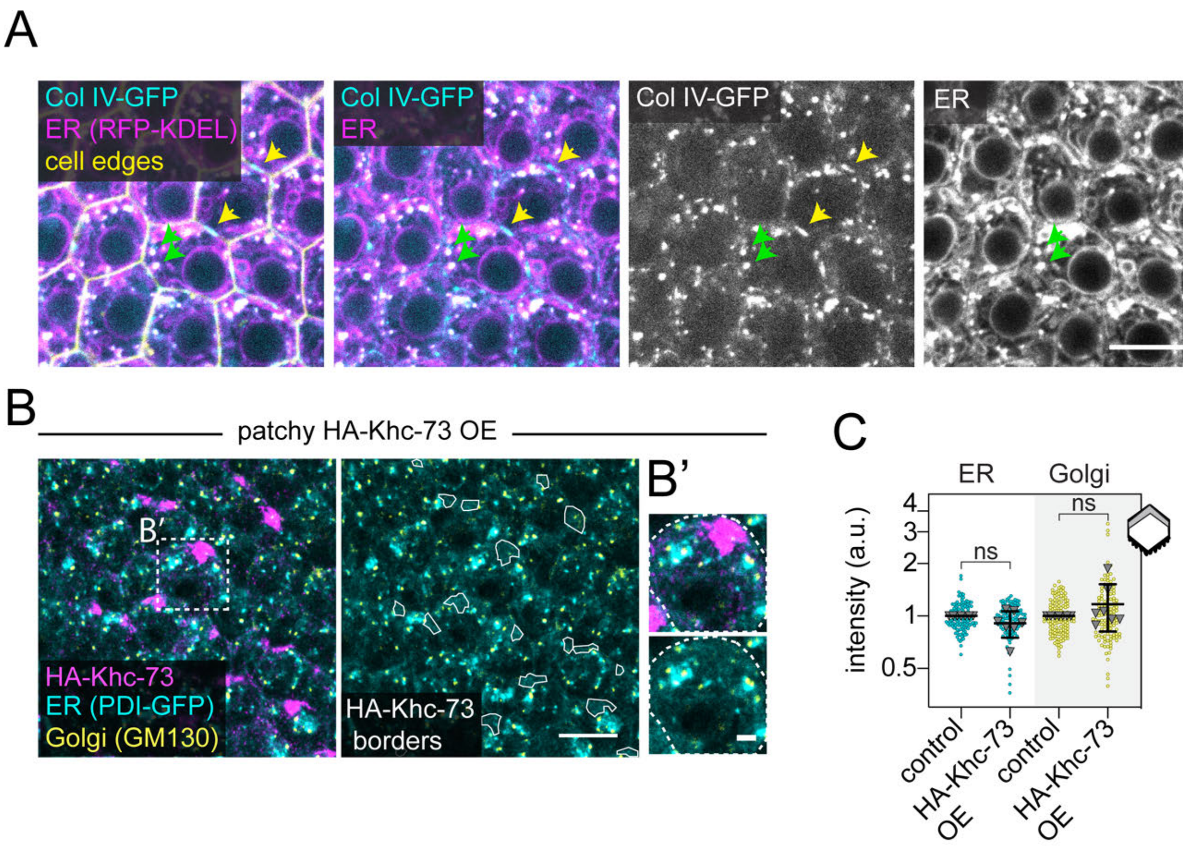
A
Col IV-GFP
ER (RFP-KDEL)
cell edges
Col IV-GFP
ER
Col IV-GFP
ER
B
patchy HA-Khc-73 OE
B′
HA-Khc-73
ER (PDI-GFP)
Golgi (GM130)
HA-Khc-73 borders
B'
C
ER
Golgi
intensity (a.u.)
4
3
2
1
0.5
ns
ns
control
HA-Khc-73 OE
control
HA-Khc-73 OE

72

**Figure S4, related to Figures 3 and 4, and Movie 3**

A. Image showing intracellular Col IV-GFP punctae colocalize with the ER luminal marker (UAS-RFP-KDEL). Cell edges are marked with CellMask. Green arrowheads highlight two Col IV-GFP punctae that overlap with 2 punctae of the ER marker. Yellow arrows point to Col IV-GFP signal that does not overlap with the ER marker, but instead with the cell edges, because it has likely been secreted.

B. Image of the ER (PDI-GFP, endogenous label) and Golgi (anti-GM130) in tissue patchily overexpressing HA-Khc-73. White lines on the right panel trace the borders of the magenta HA-Khc-73 foci from the left panel to make it easier to see the ER and Golgi labels are not enriched within these regions. B' inset highlights a single cell outlined with a dotted white line in B.

C. Quantification of the levels of the ER label and the Golgi label at the basal trailing edge of cells (grey region of cell in cartoon) that are grouped as "control" cells lacking HA-Khc-73 staining or cells overexpressing HA-Khc-73. Colored points represent individual cells and grey triangles represent the means for each egg chamber. Statistics were performed on the mean egg chamber data. Data represent mean ± SD plotted on a log scale. For ER, one sample t test compared to the theoretical ratio of 1, ns $p>0.05$. For Golgi, Wilcoxon signed rank test, ns $p>0.05$. n=7 egg chambers, 135 "control" and 121 HA-Khc-73 OE cells.

Stage 7 egg chambers. Images are oriented such that migration is towards the bottom of the page. Scale bars, 10 µm (A and B), 2 µm in B'.

### Figure S5

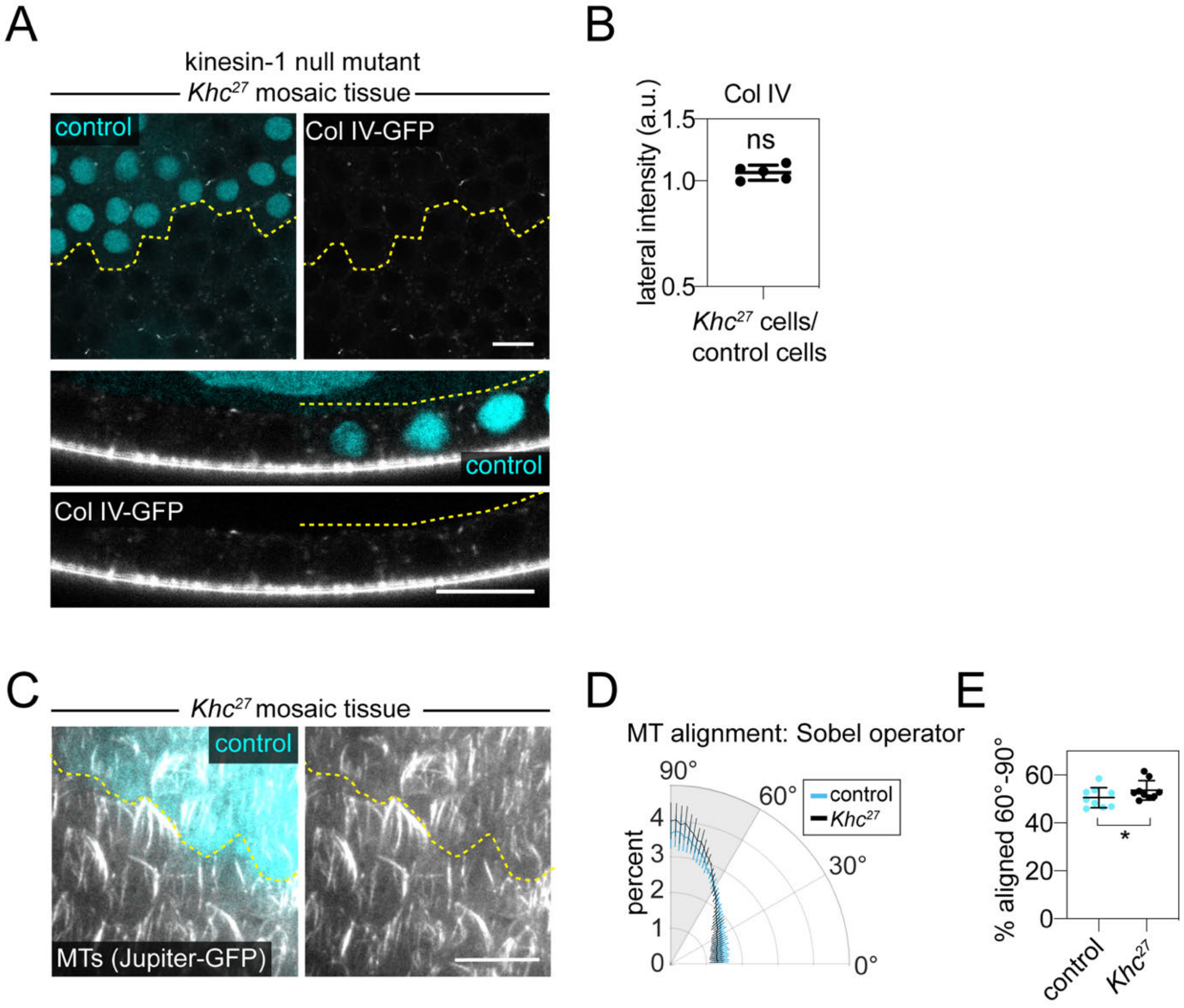
A
kinesin-1 null mutant
Khc27 mosaic tissue
control
Col IV-GFP
control
Col IV-GFP
B
Col IV
ns
lateral intensity (a.u.)
1.5
1.0
0.5
Khc27 cells/
control cells
C
Khc27 mosaic tissue
control
MTs (Jupiter-GFP)
D
MT alignment: Sobel operator
90°
60°
30°
0°
control
Khc27
percent
4
3
2
1
0
E
% aligned 60°-90°
60
40
20
0
*
control
Khc27

94

**Figure S5, related to Figures 5 and 6**

A. Images showing Col IV-GFP is localized normally in mosaic tissue of the kinesin-1 null allele $Khc^{27}$. Top panels are planes through cells' lateral domains. Bottom panels are cross sections. The dotted yellow line demarcates control and $Khc^{27}$ cells.

B. Quantification of Col IV-GFP at lateral cell edges (plane in upper panel of A) in $Khc^{27}$ cells relative to controls within the same mosaic tissue. No increase in lateral accumulation Col IV-GFP occurs in $Khc^{27}$ cells. Data are plotted on a log scale. One sample t test compared to theoretical ratio of 1, ns $p>0.05$. n=5 egg chambers.

C. Image of MTs (Jupiter-GFP, endogenously labeled MT-associated protein) from a partial TIRF time-lapse along the basal surface of a $Khc^{27}$ mosaic epithelium. MTs remain aligned along the basal surface of $Khc^{27}$ cells. The dotted yellow line demarcates control and $Khc^{27}$ cells. MT intensity appears uneven from partial TIRF illumination.

D. Quantification of MT alignment using a Sobel operator shows the degree of MT alignment in control and $Khc^{27}$ cells from (C) relative to the direction of cell migration. n=9 egg chambers. The direction of cell migration is 90º.

E. Quantification of the enrichment in MT alignment in control and $Khc^{27}$ cells. To compare how much alignment with the direction of migration occurs in both genotypes, we measured the population aligned between 60º-90º from the Sobel operator (grey shading in D), where perfect alignment with the direction of migration is 90º. $Khc^{27}$ cells have a minor, but significant, increase in aligned MTs. Wilcoxon matched-pairs signed rank test, * $p<0.05$. n=9 egg chambers.

Stage 8 egg chambers (A, B), stage 7 egg chambers (C-E). Images are oriented such that migration is toward the bottom of the page. Data represent mean ± SD in all plots. Scale bars, 10 µm.

### Figure S6

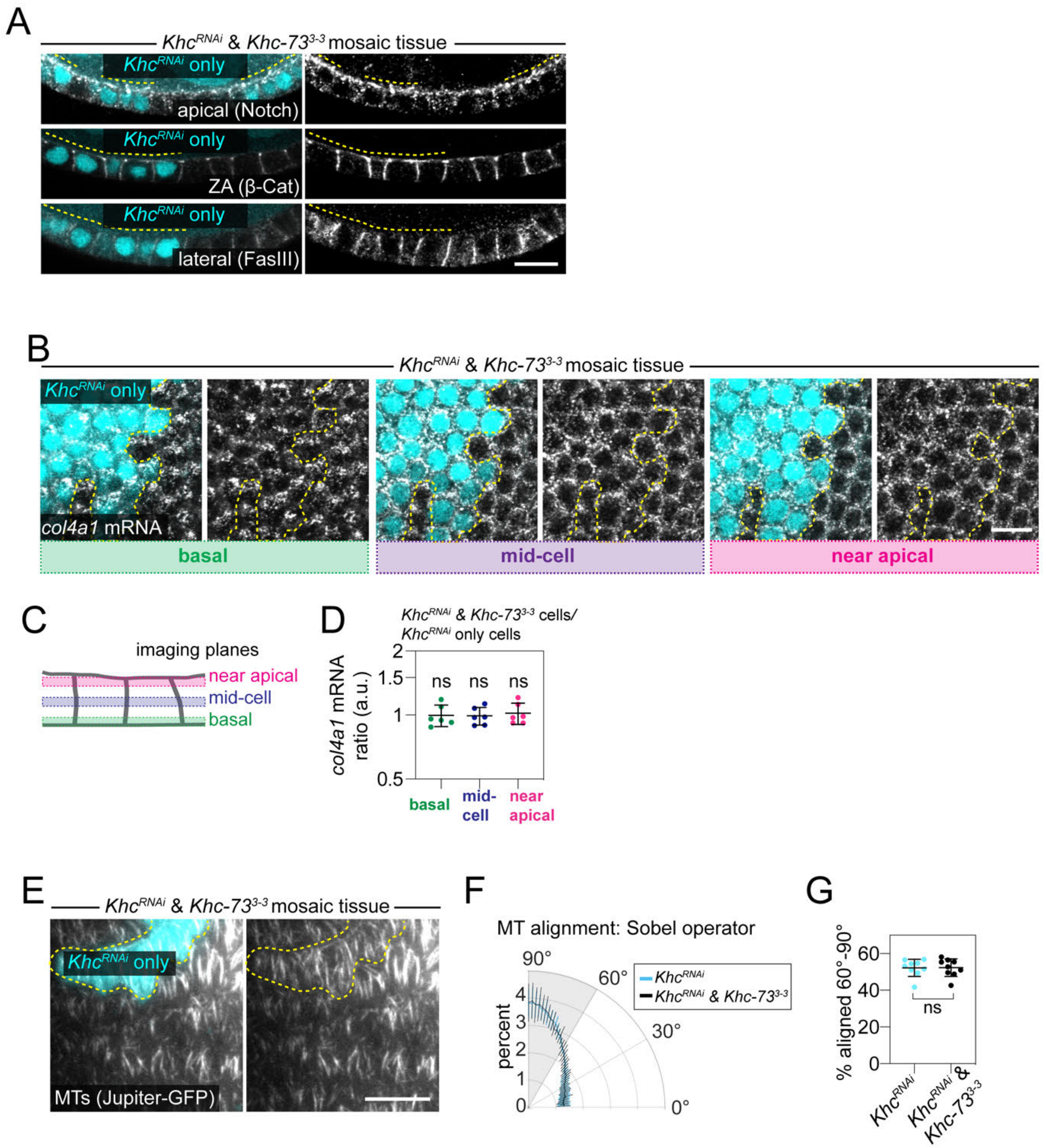

A
Khc^RNAi & Khc-73^3-3 mosaic tissue
Khc^RNAi only
apical (Notch)
ZA (β-Cat)
lateral (FasIII)
B
Khc^RNAi & Khc-73^3-3 mosaic tissue
Khc^RNAi only
col4a1 mRNA
basal
mid-cell
near apical
C
imaging planes
near apical
mid-cell
basal
D
Khc^RNAi & Khc-73^3-3 cells/
Khc^RNAi only cells
col4a1 mRNA ratio (a.u.)
ns
basal
mid-cell
near apical
E
Khc^RNAi & Khc-73^3-3 mosaic tissue
Khc^RNAi only
MTs (Jupiter-GFP)
F
MT alignment: Sobel operator
percent
Khc^RNAi
Khc^RNAi & Khc-73^3-3
G
% aligned 60°-90°
ns
Khc^RNAi
Khc^RNAi & Khc-73^3-3

**Figure S6, related to Figure 6**

A. Images showing normal localization of transmembrane proteins to apical (Notch) or lateral (FasIII) domains, and location of β-catenin to the zonula adherens in *Khc*$^{RNAi}$ & *Khc-73*$^{3\text{-}3}$ cells within a mosaic epithelium. All proteins detected by immunostaining. Cells only expressing *Khc*$^{RNAi}$ are marked in cyan and with a yellow dotted line drawn above them.

B. Images of single molecule inexpensive fluorescence in situ hybridization (smiFISH) for *col4a1* mRNA in *Khc*$^{RNAi}$ & *Khc-73*$^{3\text{-}3}$ mosaic tissue taken at 3 different z-planes through the lateral domains (see diagram in C). The dotted yellow line demarcates *Khc*$^{RNAi}$ only cells and *Khc*$^{RNAi}$ & *Khc-73*$^{3\text{-}3}$ cells.

C. Illustration of imaging planes in (B).

D. Quantification of *col4a1* mRNA intensity in *Khc*$^{RNAi}$ & *Khc-73*$^{3\text{-}3}$ cells relative to *Khc*$^{RNAi}$ only cells in the same mosaic tissue at three different z-planes. The *col4a1* mRNA distribution along the apical-basal axis does not change in *Khc*$^{RNAi}$ & *Khc-73*$^{3\text{-}3}$ cells. Data are plotted on a log scale. One sample t tests compared to the theoretical ratio of 1, ns $p>0.05$. n= 6 egg chambers.

E. Image of MTs (Jupiter-GFP, endogenously labeled MT associated protein) from a partial TIRF time-lapse along the basal surface of a *Khc*$^{RNAi}$ & *Khc-73*$^{3\text{-}3}$ mosaic epithelia. MTs remain aligned in *Khc*$^{RNAi}$ & *Khc-73*$^{3\text{-}3}$ cells. The dotted yellow line demarcates *Khc*$^{RNAi}$ only cells and *Khc*$^{RNAi}$ & *Khc-73*$^{3\text{-}3}$ cells.

F. Quantification of MT alignment using a Sobel operator shows the degree of MT alignment in *Khc*$^{RNAi}$ only cells and *Khc*$^{RNAi}$ & *Khc-73*$^{3\text{-}3}$ cells from (E) relative to the direction of cell migration. The direction of cell migration is 90º. n=9 egg chambers.

G. Quantification of the enrichment in MT alignment in *Khc*$^{RNAi}$ only cells and *Khc*$^{RNAi}$ & *Khc-73*$^{3\text{-}3}$ cells. To compare how much alignment with the direction of migration occurs in both genotypes, we measured the population aligned between 60º-90º from the Sobel operator (grey shading in F), where perfect alignment with the direction of migration is 90º. Paired t test, ns $p>0.05$. n=9 egg chambers.

Stage 7 egg chambers, except (A), which contains stages 6 and 7. Data represent mean ± SD. Scale bars, 10 μm.

**Supplemental Table 1, related to all Figures and Movies**

**Detailed experimental genotypes**

| Figure | Panel | Genotype |
|---|---|---|
| 1 | A | *w; vkg-GFP$^{CC00791}$ traffic jam-Gal4/+* |
| | D F | *w; vkg-GFP$^{CC00791}$ traffic jam-Gal4/+*<br>*w; vkg-GFP$^{CC00791}$ traffic jam-Gal4/+; UAS-spastin/+*<br>*w; vkg-GFP$^{CC00791}$ traffic jam-Gal4 Khc-73$^{3-3}$/Khc-73$^{3-3}$*<br>*rab10$^{-}$/rab10$^{-}$; vkg-GFP$^{CC00791}$/+* |
| | G | *w; traffic jam-Gal4/+; LanA-GFP$^{fTR000\ 574}$/+*<br>*w; traffic jam-Gal4/UAS-Khc-73 RNAi$^{HMS01624}$; LanA-GFP$^{fTR000\ 574}$/+*<br>*trol-GFP$^{CA06698}$/+; traffic jam-Gal4/+*<br>*trol-GFP$^{CA06698}$/+; traffic jam-Gal4/UAS-Khc-73 RNAi$^{HMS01624}$* |
| 2 | A B | *w; vkg-GFP$^{CC00791}$ traffic jam-Gal4 Khc-73$^{3-3}$/Khc-73$^{3-3}$* |
| | C E | *hsFlp/+; vkg-GFP$^{CC00791}$ traffic jam-Gal4 FRTG13 Ubi-mRFP.nls/FRTG13 Khc-73$^{3-3}$* |
| 3 | A B | *w; traffic jam-Gal4/+; UAS-YFP-Rab10/+* |
| | C | *w; traffic jam-Gal4/UAS-RFP-Rab10; Khc-73-GFP$^{fTRG1377}$* |
| | D E | *w$^{1118}$* |
| | G G' | *w; traffic jam-Gal4/UAS-ChRFP-humanαTubulin; UAS-YFP-Rab10/+* |
| 4 | A B | *hsFLP; traffic jam-Gal4 FRTG13 Ubi-mRFP.nls/FRTG13 Khc-73$^{3-3}$; UAS-YFP-Rab10/+* |
| | C D | *YFP-Rab10; traffic jam-Gal4/+; UAS-HA-Khc-73/+*<br>*w; vkg-GFP$^{CC00791}$ traffic jam-Gal4/+; UAS-HA-Khc-73/+* |
| | F G H | *w; traffic jam-Gal4/+; UAS-YFP-Rab10/+*<br>*w; traffic jam-Gal4 Khc-73$^{3-3}$/Khc-73$^{3-3}$; UAS-YFP-Rab10/+* |
| 5 | A C | *w; vkg-GFP$^{CC00791}$ traffic jam-Gal4/+; UAS-Dcr-2/+*<br>*w; vkg-GFP$^{CC00791}$ traffic jam-Gal4/+; UAS-Dcr-2/UAS-Khc RNAi$^{GL00330}$/+*<br>*w; vkg-GFP$^{CC00791}$ traffic jam-Gal4/+; UAS-Dcr-2/UAS-Klc RNAi$^{HMS00883}$/+*<br>*w; vkg-GFP$^{CC00791}$ traffic jam-Gal4/UAS-Khc-73 RNAi$^{HMS01624}$; UAS-Dcr-2/+*<br>*w; vkg-GFP$^{CC00791}$ traffic jam-Gal4/UAS-Khc-73 RNAi$^{HMS01624}$; UAS-Dcr-2/UAS-Khc RNAi$^{GL00330}$*<br>*w; vkg-GFP$^{CC00791}$ traffic jam-Gal4/UAS-Khc-73 RNAi$^{HMS01624}$; UAS-Dcr-2/UAS-Klc RNAi$^{HMS00883}$* |
| 6 | A B | *hsFLP/YFP-Rab10; FRTG13 Ubi-mRFP.nls/FRTG13 Khc$^{27}$*<br>*hsFLP/YFP-Rab10; FRTG13 Ubi-mRFP.nls/FRTG13 Khc$^{KO.mutA}$* |
| | C D | *hsFLP/+; FRTG13 Ubi-mRFP.nls/ FRTG13 Khc$^{27}$; Khc-73-GFP$^{fTRG01377}$/+* |
| | F G | *hsFLP/+; traffic jam-Gal4 FRTG13 Ubi-mRFP.nls/FRTG13 Khc-73$^{3-3}$; UAS-YFP-Rab10/UAS-Khc RNAi$^{GL00330}$* |
| | H | *w; vkg-GFP$^{CC00791}$ traffic jam-Gal4 Khc-73$^{3-3}$/Khc-73$^{3-3}$; UAS-Khc RNAi$^{GL00330}$/+* |

| | | |
|---|---|---|
| | I K | w; traffic jam-Gal4/+; UAS-YFP-Rab10/+<br>w; traffic jam-Gal4 Khc-73$^{3-3}$/Khc-73$^{3-3}$; UAS-YFP-Rab10/UAS-Khc RNAi$^{GL00330}$ |
| 7 | A | w; vkg-GFP$^{CC00791}$ traffic jam-Gal4/+; UAS-Dcr-2/+<br>w; vkg-GFP$^{CC00791}$ traffic jam-Gal4 Khc-73$^{3-3}$/Khc-73$^{3-3}$<br>w; vkg-GFP$^{CC00791}$ traffic jam-Gal4/UAS-Khc-73 RNAi$^{HMS01624}$; UAS-Dcr-2/UAS-Khc RNAi$^{GL00330}$<br>w; vkg-GFP$^{CC00791}$ traffic jam-Gal4 Khc-73$^{3-3}$/Khc-73$^{3-3}$; UAS-Khc RNAi$^{GL00330}$/+ |
| | B | w; vkg-GFP$^{CC00791}$ traffic jam-Gal4/+; UAS-Dcr-2/+<br>w; vkg-GFP$^{CC00791}$ traffic jam-Gal4/+; UAS-Dcr-2/UAS-Khc RNAi$^{GL00330}$/+<br>w; vkg-GFP$^{CC00791}$ traffic jam-Gal4/UAS-Khc-73 RNAi$^{HMS01624}$; UAS-Dcr-2/UAS-Khc RNAi$^{GL00330}$/+<br>w; vkg-GFP$^{CC00791}$ traffic jam-Gal4/+<br>w; vkg-GFP$^{CC00791}$ traffic jam-Gal4 Khc-73$^{3-3}$/Khc-73$^{3-3}$<br>w; vkg-GFP$^{CC00791}$ traffic jam-Gal4 Khc-73$^{3-3}$/Khc-73$^{3-3}$; UAS-Khc RNAi$^{GL00330}$/+ |
| | C | w; vkg-GFP$^{CC00791}$ traffic jam-Gal4/+; UAS-Dcr-2/+<br>w; vkg-GFP$^{CC00791}$ traffic jam-Gal4 Khc-73$^{3-3}$/Khc-73$^{3-3}$<br>w; vkg-GFP$^{CC00791}$ traffic jam-Gal4/UAS-Khc-73 RNAi$^{HMS01624}$; UAS-Dcr-2/UAS-Khc RNAi$^{GL00330}$/+<br>w; vkg-GFP$^{CC00791}$ traffic jam-Gal4 Khc-73$^{3-3}$/Khc-73$^{3-3}$; UAS-Khc RNAi$^{GL00330}$/+ |
| | D | w; vkg-GFP$^{CC00791}$ traffic jam-Gal4/+; UAS-Dcr-2/+<br>w; vkg-GFP$^{CC00791}$ traffic jam-Gal4/+; UAS-Dcr-2/UAS-Khc RNAi$^{GL00330}$/+<br>w; vkg-GFP$^{CC00791}$ traffic jam-Gal4/UAS-Khc-73 RNAi$^{HMS01624}$; UAS-Dcr-2/UAS-Khc RNAi$^{GL00330}$/+<br>w; vkg-GFP$^{CC00791}$ traffic jam-Gal4/+<br>w; vkg-GFP$^{CC00791}$ traffic jam-Gal4 Khc-73$^{3-3}$/Khc-73$^{3-3}$ |
| | E | w; vkg-GFP$^{CC00791}$ traffic jam-Gal4/+<br>w; vkg-GFP$^{CC00791}$ traffic jam-Gal4 Khc-73$^{3-3}$/Khc-73$^{3-3}$ |
| S1 | B | w; vkg-GFP$^{CC00791}$ traffic jam-Gal4/+; UAS-spastin/+ |
| | D E | w; vkg-GFP$^{CC00791}$ traffic jam-Gal4 Khc-73$^{3-3}$/ +<br>w; vkg-GFP$^{CC00791}$ traffic jam-Gal4 Khc-73$^{3-3}$/Khc-73$^{149}$ |
| | F | w; vkg-GFP$^{CC00791}$ traffic jam-Gal4 Khc-73$^{3-3}$/ Khc-73$^{3-3}$ |
| S2 | A B D | hsFlp/+; FRTG13 Ubi-mRFP.nls/FRTG13 Khc-73$^{3-3}$<br>hsFlp/+; vkg-GFP$^{CC00791}$ traffic jam-Gal4 FRTG13 Ubi-mRFP.nls/FRTG13 Khc-73$^{3-3}$<br>A mixture of these two genotypes was used, differing only in expression of 1 copy of Col IV-GFP (endogenous) and *traffic jam-GAL4* (with no UAS transgenes expressed) |
| S3 | A-E | hsFLP/+; FRTG13 Ubi-mRFP.nls/FRTG13 Khc-73$^{3-3}$; Jupiter-GFP$^{G00147}$/+ |
| | F G | hsFLP; FRTG13 Ubi-mRFP.nls/FRTG13 Khc-73$^{3-3}$; Ubi-PatroninA-GFP/+ |

| | | |
|---|---|---|
| S4 | A | w; *vkg-GFP*$^{CC00791}$ *traffic jam-Gal4/+; UAS-RFP-KDEL/+* |
| | B C | *w; traffic jam-Gal4/+; PDI-GFP*$^{CPTI000688}$*/UAS-HA-Khc-73* |
| S5 | A B | *hsFLP/+; FRTG13 Khc*$^{27}$*/vkg-GFP*$^{CC00791}$ *traffic-jam-Gal4 FRTG13 Ubi-mRFP.nls* |
| | C D F | *hsFLP/+; FRTG13 Khc*$^{27}$*/FRTG13 Ubi-mRFP.nls; Jupiter-GFP*$^{G00147}$*/+* |
| S6 | A B D | *hsFLP/+; FRTG13 Khc-73*$^{3\text{-}3}$*/vkg-GFP*$^{CC00791}$ *traffic jam-Gal4 FRTG13 Ubi-mRFP.nls; UAS-Khc RNAi*$^{GL00330}$*/+* |
| | E F G | *hsFLP/+; FRTG13 Khc-73*$^{3\text{-}3}$*/traffic jam-Gal4 FRTG13 Ubi-mRFP.nls; UAS-Khc RNAi*$^{GL00330}$*/Jupiter-GFP*$^{G00147}$ |
| Movie 1 | | *w; vkg-GFP*$^{CC00791}$ *traffic jam-Gal4* |
| Movie 2 | | *w1118* |
| Movie 3 | | *w; traffic jam-Gal4/UAS-ChRFP-human*$\alpha$*Tubulin; UAS-YFP-Rab10/+* |
| Movie 4 | | *w; traffic jam-Gal4/+; UAS-YFP-Rab10/UAS-HA-Khc-73* |
| Movie 5 | | *w; traffic jam-Gal4/+; UAS-YFP-Rab10/+*<br>*w; traffic jam-Gal4 Khc-73*$^{3\text{-}3}$*/Khc-73*$^{3\text{-}3}$*; UAS-YFP-Rab10/+* |
| Movie 6 | | *w; vkg-GFP*$^{CC00791}$ *traffic jam-Gal4/+*<br>*w; vkg-GFP*$^{CC00791}$ *traffic jam-Gal4 Khc-73*$^{3\text{-}3}$*/Khc-73*$^{3\text{-}3}$ |

**Supplemental Table 2, related to all Figures and Movies**

**Experimental conditions**

Adult females were selected 1-2 days after eclosion for experiments, and yeasted in the presence of males. Temperature and number of days females were on yeast is detailed for each experiment. If heat-shock was done to induce Flp-FRT mediated recombination prior to yeasting, it is noted as “heat-shock + 25°C”. For all heat-shock experiments, adult flies were cycled through the following heat-shock protocol in an ECHOtherm™ chilling incubator for 2 weeks: 1 hr 37ºC, 1 hr 25ºC, 1 hr 37ºC, every 12 hours, holding at 25°C in between. Flies were transferred to new vials every 4 days. The final 2 days, flies were moved to fresh vials on yeast.

| Figure | Panels | Temperature | # days ♀ on yeast |
|---|---|---|---|
| 1 | A | 25°C | 2 days |
| | D F | 25°C | 1 day, *rab10*$^{-}$<br>2 days, all other genotypes |
| | G | 29°C | 3 days |
| 2 | A B | 25°C | 2 days |
| | C E | heat-shock + 25°C | 2 days |
| 3 | A-G | 25°C | 2 days |
| 4 | A B | heat-shock + 25°C | 2 days |
| | C-H | 25°C | 2 days |
| 5 | A C | 29°C | 3 days |
| 6 | A-G | heat-shock + 25°C | 2 days |
| | H-K | 25°C | 1 day |
| 7 | A-E | 29°C | 3 days, all RNAi panels and their control |
| | | 25°C | 2 days, *Khc-73*$^{3-3}$ and its control |
| | | 25°C | 1 day, *Khc*$^{RNAi}$ & *Khc-73*$^{3-3}$ |
| S1 | B, D-F | 25°C | 2 days |
| S2 | A-D | heat-shock + 25°C | 2 days |
| S3 | A-G | heat-shock + 25°C | 2 days |
| S4 | A-C | 25°C | 2 days |
| S5 | A-E | heat-shock + 25°C | 2 days |
| S6 | A-G | heat-shock + 25°C | 2 days |
| Movie 1 | | 25°C | 2 days |
| Movie 2 | | 25°C | 2 days |
| Movie 3 | | 25°C | 2 days |
| Movie 4 | | 25°C | 2 days |
| Movie 5 | | 25°C | 2 days |
| Movie 6 | | 25°C | 2 days |
